# Correlative light–electron microscopy reveals fundamentally different axonal architectures for fast and modulatory transmission

**DOI:** 10.64898/2026.08.11.744243

**Authors:** KS Lee, DA Sahlender, JR Blanc, A Jorstad, C Gasselin, S Clerc-Rosset, F Markopoulos, CCH Petersen, A Holtmaat, GW Knott

**Affiliations:** Bio Electron Microscopy Facility, School of Life Sciences, EPFL, Lausanne, Switzerland; Laboratory of Sensory Processing, School of Life Sciences, EPFL, Lausanne; Department of Basic Neurosciences, University of Geneva, Switzerland; Computer Vision Laboratory, School of Computer and Communication Sciences, EPFL, Switzerland

## Abstract

We used correlative light and focussed ion beam–scanning electron microscopy (FIBSEM) to characterise the ultrastructural organisation of neuromodulatory axons in the upper layers of the adult mouse somatosensory cortex. Cholinergic projections from the basal forebrain and serotonergic axons from the raphe nucleus were reconstructed, enabling quantitative analysis of synapse and vesicle distribution. Consistent with prior work, these axons formed only small and morphologically indistinct synaptic specialisations. However, these sites contained only a small fraction of the total axonal vesicle population, which were instead distributed along their length and spatially uncoupled from identifiable synaptic contacts. In contrast to classical excitatory and inhibitory axons, where vesicles are tightly clustered at defined presynaptic release sites, this organisation indicates that synaptic release contributes only minimally to neuromodulator output. Instead, our data support a model in which acetylcholine and serotonin signalling is mediated predominantly through non-synaptic release, consistent with a spatially diffuse transmission across cortical microcircuits.

## Introduction

Neural communication is classically understood to occur at synapses, specialised cell junctions, where presynaptic regions containing clusters of vesicles are apposed to a postsynaptic density, enabling fast and spatially precise signaling. This view of synaptic organisation first emerged from early transmission electron microscopy studies in the mid-twentieth century. These established ultrastructural criteria for synapse identification in the brain, including vesicle accumulation, pre and postsynaptic membrane specialisations (Gray, 1959a, 1959b; Palay, 1956; Peters et al., 1991).

Today, however, not all neurotransmitter systems are thought to conform to the classical model of point-to-point synaptic transmission. Neuromodulatory transmitters such as acetylcholine and serotonin exhibit properties suggestive of alternative modes of signalling (Descarries & Mechawar, 2000). Anatomical evidence using immunoelectron microscopy to identify serotonergic and cholinergic axons has shown that synapses of these neuromodulatory axons are comparatively sparse, small, or morphologically ambiguous. These studies consistently report a high proportion of axonal varicosities lacking clear postsynaptic densities and other classical synaptic specialisations (Beaudet & Descarries, 1976; Descarries et al., 1975; Smiley & Goldman-Rakic, 1996; Takács et al., 2018; Turrini et al., 2001; Umbriaco et al., 1994). Nevertheless, physiological evidence indicates that both acetylcholine and serotonin are capable of mediating rapid, temporally precise synaptic transmission. Fast serotonergic postsynaptic responses mediated by ionotropic 5-HT3 receptors have been recorded in neocortical interneurons, demonstrating direct communication by serotonergic afferents (Férézou et al., 2002). Likewise, optogenetic activation of basal forebrain cholinergic projections evokes short-latency nicotinic postsynaptic currents in cortical interneurons, providing evidence for fast, point-to-point cholinergic signalling (Bennett et al., 2012). Together, these findings demonstrate that neuromodulatory systems retain the capacity for conventional synaptic communication despite the apparent scarcity of morphologically identifiable synapses revealed in the ultrastructural studies. Yet despite the morphological and physiological evidence for both synaptic and extrasynaptic modes of signalling by acetylcholine and serotonin, the structural organisation of these axons remains poorly understood. Previous ultrastructural studies have focussed primarily on individual varicosities and whether they possess the hallmarks of synaptic release sites. While this approach has established that many neuromodulatory varicosities lack synapses, it provides only a limited view of axonal organisation. Connectivity is primarily inferred from the presence or absence of membrane specialisations at the individual varicosities that are targeted by electron microscopy, while little attention is paid to the overall distribution of putative synaptic contacts along the axon or to the location of the neurotransmitter-containing vesicles. Consequently, it remains unclear how neuromodulatory axons are organised relative to the fast transmitting systems, whether vesicles are predominantly associated with structurally discernible synaptic contacts, and to what extent synapses represent the principal sites of transmitter release. Therefore, a structural analysis that considers the entire axon, rather than individual varicosities in isolation, is therefore required to understand the relationship between connectivity, vesicle organisation, and modes of neurotransmission in these systems.

To achieve this, we used a correlative light and electron microscopy approach, combining molecular identification of acetylcholine and serotonin axons with focussed ion beam scanning electron microscopy (FIBSEM) in mouse somatosensory cortex. Targeting modulatory axons using genetically encoded fluorescent markers overcame the limitations of electron-dense labelling strategies that obscure the axons’ contents and enabled detailed three-dimensional reconstructions of extended axonal segments. These reconstructions revealed that although neuromodulatory axons form small and often indistinct putative synaptic contacts, the vast majority of vesicles are not associated with these sites. Rather than representing a sparsely connected version of the classical synaptic network, acetylcholine and serotonin axons exhibit a pattern of vesicle organisation that differs fundamentally from that of fast transmitting axons, consistent with a mode of communication that is diffuse, non-specific, and only weakly tied to individual postsynaptic partners.

## Results

### Correlative identification and reconstruction of molecularly-defined axons

We have previously established a workflow that enables reliable identification and three-dimensional reconstruction of defined axons and dendrites in the adult mouse brain (Maco et al., 2014). In the current study, acetylcholine and serotonin axons were genetically labeled via Cre-dependent viral expression of fluorescent reporters in basal forebrain cholinergic neurons (ChAT-Cre mice) and raphe serotonergic neurons (SERT-Cre mice). This produced robust axonal fluorescence in the somatosensory cortex (Figure 1A).

**Figure 1.**
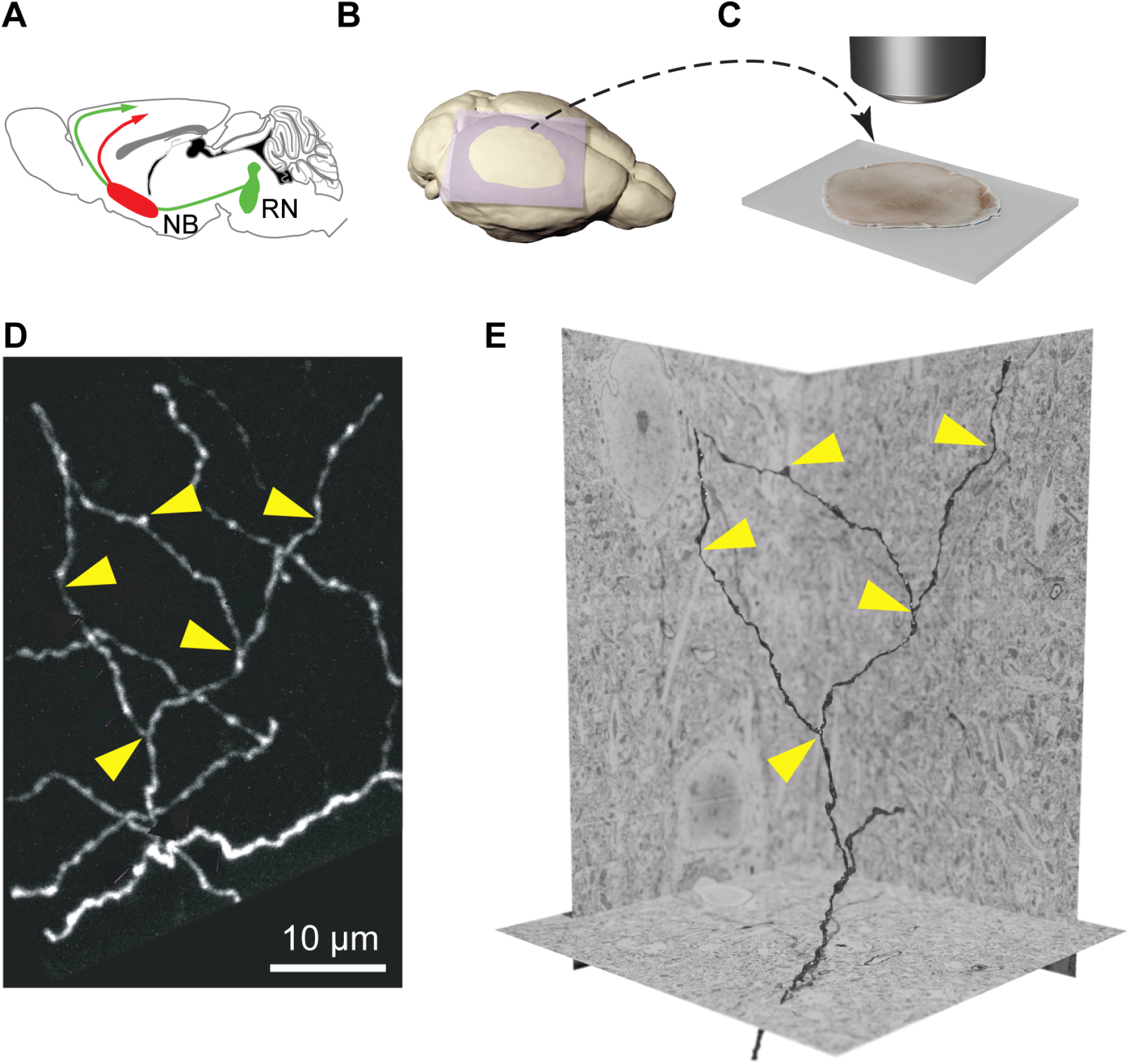
Correlative light and FIBSEM of fluorescent axons. **(A)** Acetylcholine and serotonin axons projecting to the somatosensory cortex were labelled via viral injections into the nucleus basalis (NB), and raphe nucleus (RN), and imaged through cranial windows to check the quality of the labelling. Cholinergic axons were labeled using AAV-FLEX_GCaMP6s-mRuby virus injections in adult mice positive for ChAT-Cre genotyping, and serotonergic axons were labelled using AAV-CAG-Flex-tdTomato-WPRE-bGH virus injections in adult mice positive for SERT-Cre genotyping (see Methods for details). **(B)** After chemical fixation, the brain was removed and vibratome sections cut tangentially to the somatosensory cortex. **(C)** These sections were viewed with fluorescent microscopy and axons of interest selected for FIBSEM imaging. **(D)** Volumes of tissue were imaged, and axons corresponding to those imaged with fluorescent microscopy were segmented on each image place that resulted in a 3D model **(E)**.

The fluorescent acetylcholine and serotonin axons were visible in brain sections, after fixation in buffered aldehyde that optimally preserved the ultrastructure. Therefore, by using fluorescence imaging, prior to heavy metal staining, and resin embedding, we could map their precise position relative to fiducial marks, such as blood vessels, and laser marks (see methods, Figure 1), so that the exact volume in which they were situated could be imaged with the FIBSEM (Figure 2A), yielding high-resolution volumetric datasets in which the axons could be unambiguously identified, segmented, and reconstructed (Figure 2B, C, D). This method avoids using any electron dense staining of the axons of interest as this can obscure ultrastructural details and interfere with the detailed reconstructions. Here we could make complete and accurate three-dimensional models of individual acetylcholine and serotonin releasing axons that could be quantitatively compared with those of glutamate and GABA containing axons. These reconstructions provided the structural basis for all subsequent quantitative analyses of axonal geometry, subcellular organization of vesicles, synapses and mitochondria, and their arrangement in the surrounding neuropil.

**Figure 2.**
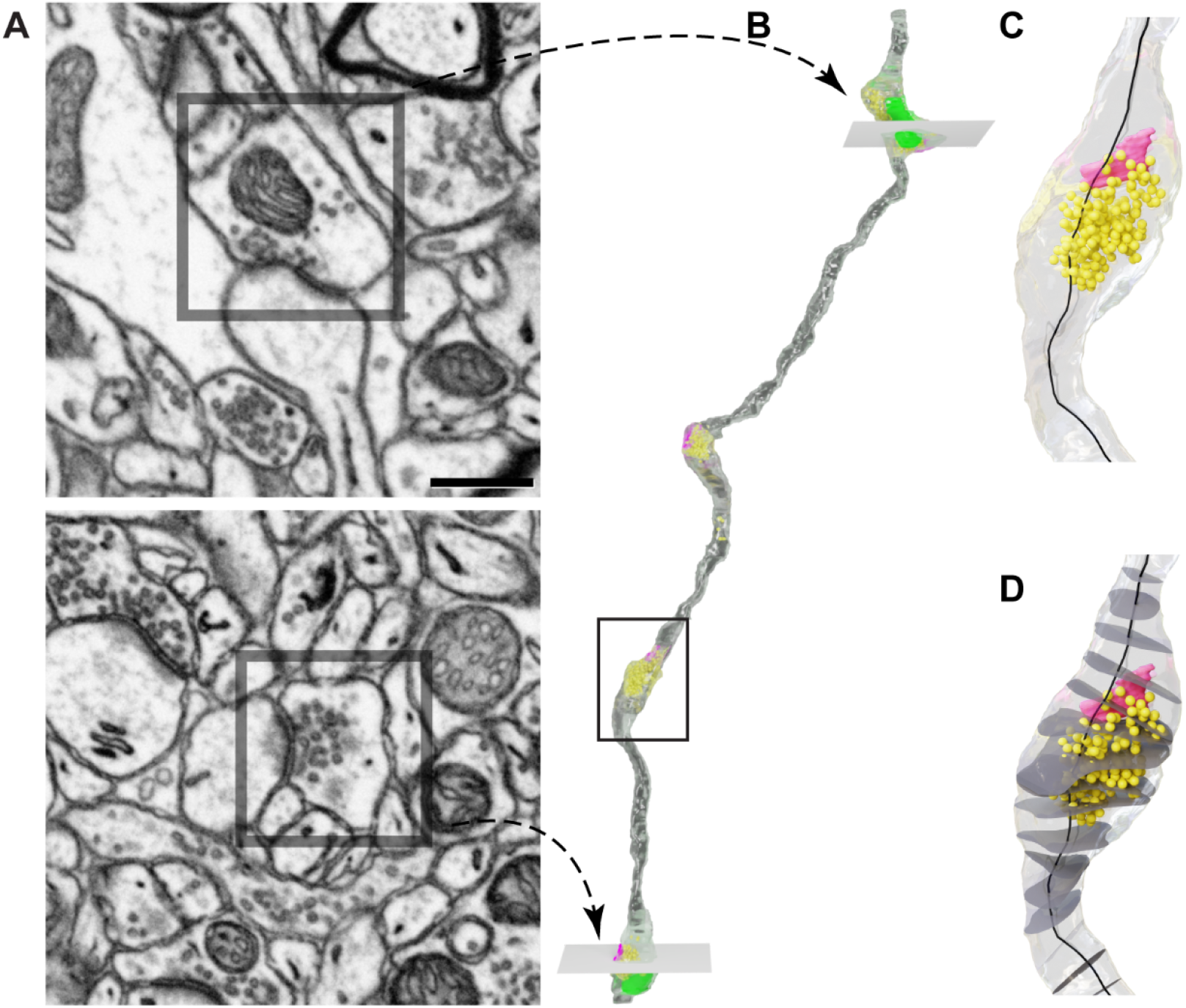
Morphometric measurements from 3D models. **(A)** Two scanning electron micrographs showing two varicosities (grey boxes) from a single glutamate axon imaged with FIBSEM. Each varicosity shows a synaptic connection and its position along the axon is indicated with a dotted arrow on the reconstruction shown in **(B)**. **(B)** reconstructed glutamate axon (grey) with vesicles (yellow), mitochondria (green) and synaptic connections (pink). The position and orientation of the electron micrographs in **(A)** are indicated with arrows, by two square grey planes. **(C)** A closer view of the axon shows a single varicosity with all the vesicles (yellow) and the synapse (pink). Also shown is a black centreline, running the length of the axon **(D)**. Cross-sectional planes (dark grey) positioned perpendicular to the black centreline at regular intervals (100 nm in our analysis) illustrate the approach used for the vesicle quantification. Rather than analysing vesicle numbers within individual varicosities, they were quantified along the entire axon by counting them within each consecutive 100 nm segment defined along the centreline.

While the acetylcholine and serotonin axons were selected using this correlative fluorescence and FIBSEM imaging, glutamate and GABA axons were identified in the same serial image stacks as the neuromodulatory axons. Glutamate axons were identified by the presence of asymmetric synapses with prominent postsynaptic densities (see Figure 2A), while GABA axons were identified by the presence of symmetric synapses with thinner postsynaptic densities and characteristic vesicle morphology (see Figure 3D). This is consistent with classical electron microscopy descriptions (Gray, 1959b; Peters et al., 1991; Uchizono, 1965). Using these methods of identification, we classified these axons accordingly: glutamate, GABA, acetylcholine, and serotonin.

**Figure 3.**
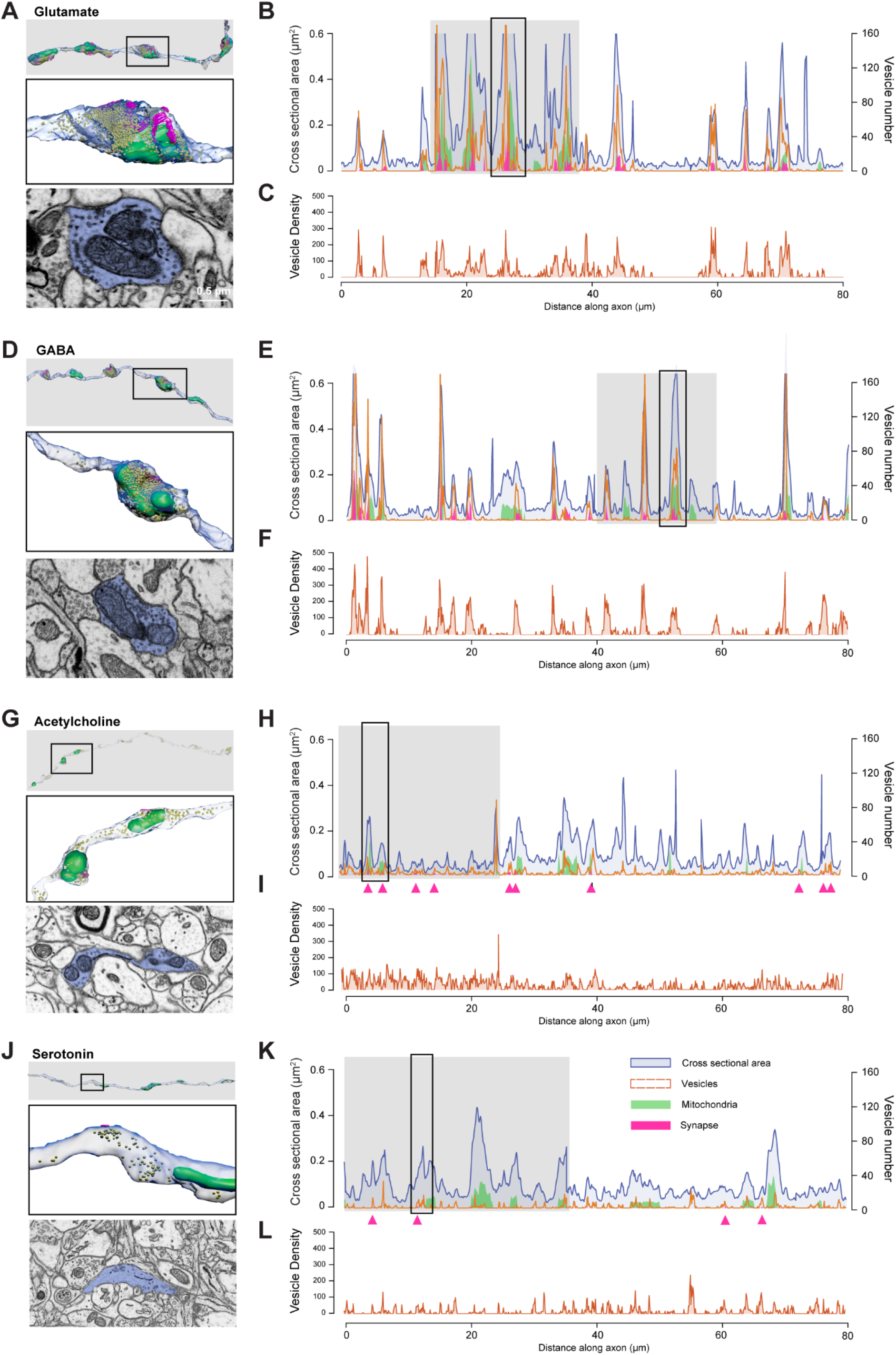
Axonal geometry and organelle distribution along the centreline. Left-hand images show 3D reconstructions of segments of glutamate **(A)**, GABA **(D)**, acetylcholine **(G)** and serotonin **(J)** axons with the solid black square indicating the higher magnification view in the image underneath of a single varicosity, or axon swelling. Below each reconstruction is a single micrograph from the series of images used to make the model, with the axon pseudocoloured in blue. The reconstructions show the vesicles (yellow), mitochondria (green) and synaptic contacts (pink). The righthand graphs **(B, E, H, K)** show the change in cross-sectional area of the axon (blue) and its mitochondria (green), plotted on the left x-axis, and vesicle number (orange) on the right y-axis for each class of axon. The grey box highlights the axon segment shown in the upper left panel of the figure. Pink arrowheads on the x-axis **(graphs H and K)** indicate the positions of synapses. The lower graphs **(C, F, I, L)** show the change in the vesicle density along the axons per µm2 of cross-sectional area.

Fluorescence light microscopy of the acetylcholine and serotonin axons did not show distinct boutons (Figure 1C, shows an acetylcholine axon), characteristic of the fast neurotransmitters, glutamate and GABA, that indicate the predominant sites of synaptic connections (Gala et al., 2017). The images, therefore, gave little indication as to where the synaptic contacts might be found. This was confirmed by the resulting 3D reconstructions, in which boutons or varicosities could not be reliably identified. For this reason, boutons and varicosities were not used as the analytical unit in this study.

Instead, a centreline was plotted through each reconstructed axon, and cross-sectional planes were generated at 100 nm intervals perpendicular to this centreline along its entire length. These cross-sections provided a continuous framework for all subsequent analyses. The cross-sectional area at each interval was used to describe axonal morphology. The same cross-sectional planes were also used to quantify the number of vesicles, as well as the extent of synaptic and mitochondria areas, at corresponding positions along the axon (see Figure 3B, E, H, K). This approach enabled us to relate local changes in axonal morphology to the spatial distribution of internal axonal structures.

Observing the profiles of the different features: cross-sectional surface area, vesicle number, axon surface area, mitochondrial surface area, indicated substantial heterogeneity, with large fluctuations in these parameters (Figure 3). In glutamate and GABA axons (Figure 3A-F), large increases in cross-sectional area coincide with peaks in vesicle number. These appear to correspond to the presence of synapses, identified using conventional ultrastructural features of presynaptic active zones, with clustered vesicles apposed to a postsynaptic densities (representative examples are shown in Figure 2 - supplemental figure 1). This is clearly evident when vesicle density is plotted per unit cross-sectional area (Figure 3C, F) where large peaks are interspersed with flat stretches with no vesicles at all. Often, but not always, these high concentrations of vesicles were accompanied by the presence of mitochondria. In the neuromodulatory types, acetylcholine and serotonin (Figure 3G-L), this correspondence was far less apparent. The changes in the axon cross-sectional area were less pronounced and the fluctuations in vesicle number were less exaggerated (Figure 3H, K), with fewer, and shorter stretches, without vesicles. Regions with increased numbers of vesicles do not appear to show any sort of indication of synapse and mitochondrial presence.

The qualitative differences in the distributions of cross-sectional area across transmitter types (Figure 4A) were further evaluated by testing for variability differences between neurotransmitter types. Given the non-normal distribution, an omnibus Levene’s test centered at the median (Brown–Forsythe design) was used. This analysis revealed a highly significant difference in structural dispersion among the groups (*W* = 75.75, *p* = 4.43 × 10⁻⁴⁸). Descriptive statistics indicated a clear hierarchy of variability: glutamate axons exhibited the highest variance (σ^2^ = 0.03) and standard deviation (SD = 0.17), followed by GABAergic axons (σ^2^ = 0.01, SD = 0.12). Acetylcholine (σ^2^ = 0.005, SD = 0.07) and serotonin (5HT; σ^2^ = 0.005, SD = 0.07) populations demonstrated the lowest variance for their axon calibre.

**Figure 4.**
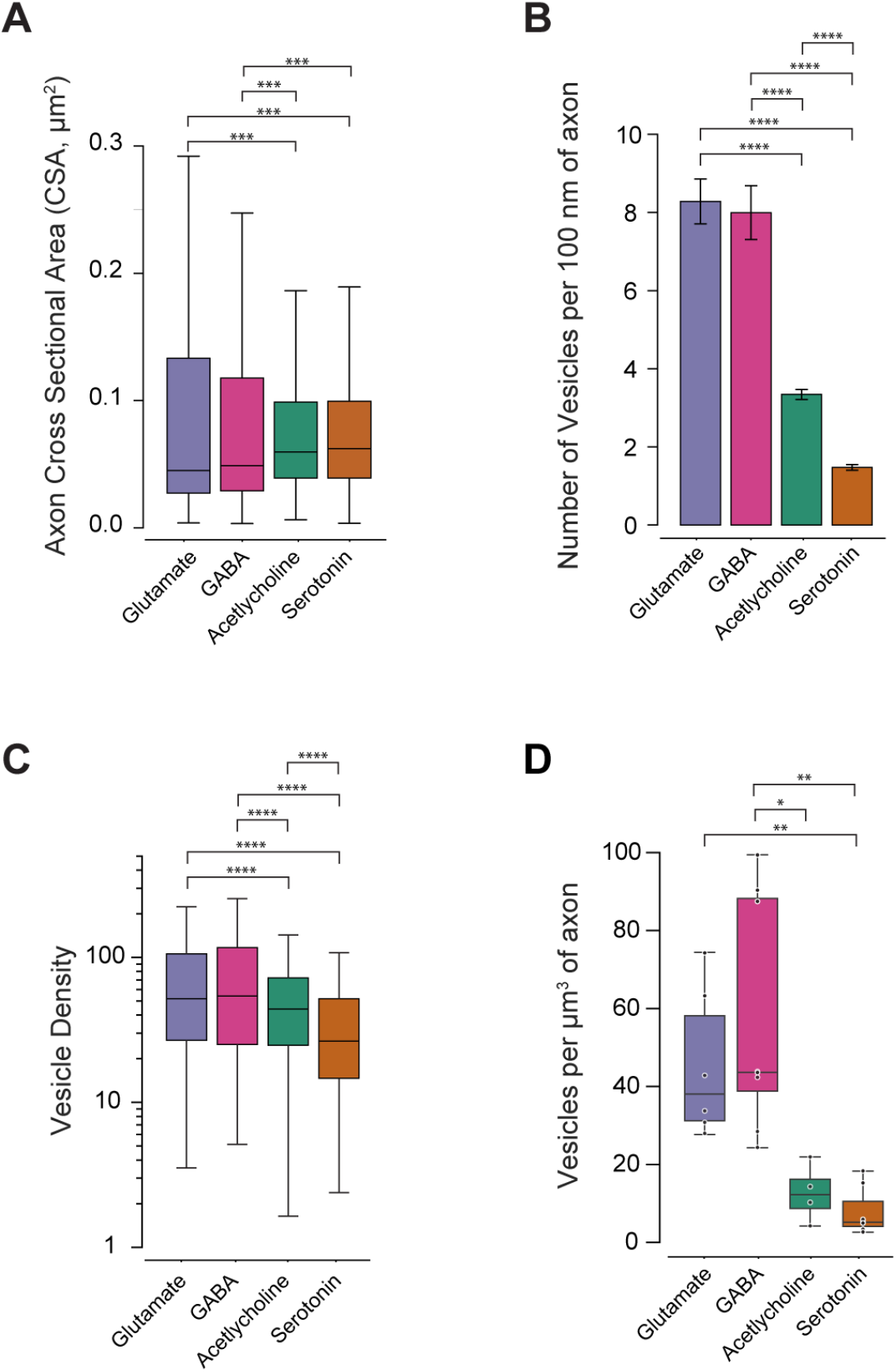
Cross-sectional area of the four types of axon, and their densities of vesicles. **(A)** Whisker box plot of cross-sectional area at every 100 nm interval along the four different types of axons; median, [Q1, Q3], n = number of slices. Glutamate: 0.0451, [0.0273, 0.1333], n = 1187, GABA: 0.0489, [0.0291, 0.1177], n = 1059; Acetylcholine: 0.0597, [0.0391, 0.0988], n = 1646; serotonin: 0.0623, [0.0391, 0.0995], n = 2076. Statistical comparisons were made with Mann–Whitney U tests, with Bonferroni correction for multiple comparisons, Glutamate vs Acetylcholine *U* = 891036, *p* = 3.84 x 10^-4^, Glutamate vs Serotonin *U* = 1125655, *p* = 2.36 x 10^-4^, GABA vs Acetylcholine *U* = 793547, *p* = 5.00 x 10^-4^, GABA vs Serotonin *U* = 996816, *p* = 1.16 x 10^-4^. **(B)** Mean number of vesicles in each 100 nm segment of glutamate, GABA, acetylcholine, and serotonin axon. Every segment was included in the analysis, even those segments containing no vesicles, therefore, mean ± SEM are shown as a high number of zero values (segments with no vesicles) were in the dataset. Glutamate: 8.28 ± 0.57, n = 1187, GABA: 8.00 ± 0.69, n = 1058, acetylcholine: 3.34 ± 0.13, n = 1646 and serotonin 1.47 ± 0.07, n = 2076; One-way ANOVA between all four neurotransmitters, F-statistic 95.51 *p* = 2.59 x 10^-60^. Welch’s t-test with Bonferroni’s correction, glutamate to acetylcholine, *p* = 7.89 x 10^-16^; glutamate to serotonin, *p* = 1.69 x 10^-29^; GABA to (continued from B) acetylcholine *p* = 3.05 x 10-^10^; GABA to serotonin p = 1.78 x 10-^19^, acetylcholine to serotonin p = 3.46 x 10^-35^ **(C)** Vesicles density per unit cross-sectional area, only segments containing vesicles were included in the dataset; median [Q1, Q3] n = number of 100 nm segments, Glutamate: 51.93 [26.84, 105.63] µm^-2^, n = 566; GABA: 54.11 [25.10, 116.93] µm^-2^ n = 450, acetylcholine: 44.10 [24.73, 72.15] µm^-2^, n = 1117, and serotonin: 26.42 [14.64, 51.89] µm^-2^, n = 788. Mann–Whitney U test, glutamate to acetylcholine *U* = 362849, p = 4.10 x 10^-6^; glutamate to serotonin *U* = 307428, p = 7.42 x 10^-32^; GABA to acetylcholine *U* = 290705, *p* = 7.08 x 10^-6^; GABA to serotonin *U* = 244570, *p* = 6.19 x 10^-28^; acetylcholine to serotonin *U* = 565775, *p* = 1.31 x 10^-25^. **(D)** Vesicles per unit volume - unit length of axon. Median [Q1, Q3] n = number of axon segments, Glutamate: 38.07 [31.22, 58.17] µm^-3^, n = 6; GABA: 43.65 [38.81, 88.24] µm^-3^, n = 8; acetylcholine: 12.25 [8.71, 16.20] µm^-3^, n = 4; serotonin: 5.15 [4.10, 10.51] µm^-3^, n = 7; Mann–Whitney U test, glutamate to serotonin *U* = 42, *p* = 7.00 x 10^-3^; GABA to acetylcholine *U* = 32, *p* = 2.42 x 10^-2^; GABA to serotonin *U* = 56, *p* = 1.86 x 10^-3^.

Turning to the contents of the axons, as the profile plots (Figure 3) show that fast transmitting axons, glutamate and GABA, have large concentrations of vesicles in regions where the axons swell. We therefore examined how these are distributed along the axon segments and whether the pattern of vesicle packing differs between the fast-transmitting and the neuromodulatory axons. If we compare the average number of vesicles per 100 nm segment, including the segments where there were none, we found that neuromodulatory axons contained, per 100 nm of axon length, far fewer vesicles than glutamate and GABA (Figure 4B; shows mean ± SEM; as this dataset contained a high proportion of zero values, and boxplot representations would not be informative, therefore a bar graph is shown; Glutamate: 8.28 ± 0.57, *n* = 1187; GABA: 8.00 ± 0.69, *n* = 1058; acetylcholine: 3.34 ± 0.13, *n* = 1646; serotonin 1.47 ± 0.07, *n* = 2076; one-way ANOVA between all four neurotransmitters, *F*-statistic 95.51 *p* = 2.59 x 10^-60^. Welch’s *t-test* with Bonferroni’s correction, glutamate to acetylcholine, *p* = 7.89 x 10^-16^; glutamate to serotonin, *p* = 1.69 x 10^-29^; GABA to acetylcholine *p* = 3.05 x 10^-10^; GABA to serotonin *p* = 1.78 x 10^-19^, acetylcholine to serotonin *p* = 3.46 x 10^-35^).

If we normalise this number to the cross-sectional area of the axon, removing the zero values, the acetylcholine and serotonin axons still showed significantly less vesicles, than glutamate and GABA (Figure 4C, median [Q1, Q3], Glutamate: 51.93 [26.84, 105.63] µm^-2^, *n* = 566, GABA: 54.11 [25.10, 116.93] µm^-2^, *n* = 450, acetylcholine: 44.10 [24.73, 72.15] µm^-2^, *n* = 1117, and serotonin: 26.42 [14.64, 51.89] µm^-2^, *n* = 788; Mann–Whitney U test, glutamate to acetylcholine *U* = 362849, *p* < 0.0001; glutamate to serotonin *U* = 307428, *p* < 0.0001; GABA to acetylcholine *U* = 290705, *p* < 0.0001; GABA to serotonin *U* = 244570, *p* < 0.0001;). There is also a significant difference between the two neuromodulatory axons, with acetylcholine containing more vesicles than serotonin (Figure 4C; acetylcholine: 44.10 [24.73, 72.15] µm^-2^, *n* = 1117, and serotonin: 26.42 [14.64, 51.89] µm^-2^; Mann–Whitney U test, acetylcholine to serotonin *U* = 565775, *p* < 0.0001).

To further compare vesicle packing within the axons, we normalised the number of vesicles to the total reconstructed volume, calculating the number of vesicles per µm^3^ per unit length (Figure 4D) for each individual axon segment. Consistent with the analysis of cross-sectional area packing density (Figure 4C), glutamate and GABA axons contained higher vesicle densities than acetylcholine and serotonin, indicating that the greater vesicle packing in the fast transmitting axons is maintained when expressing vesicle content relative to the total axon volume.

To determine whether vesicle content scales with axon calibre across neuronal subtypes, we examined the allometric relationship between vesicle number and axon cross-sectional area using log–log regression to estimate the power-law exponent. While we observed a significant positive correlation between cross-sectional area and vesicle number across all four subtypes (*p* < 0.001), the nature of this scaling varied considerably. Glutamatergic and GABAergic axons exhibited nearly isometric scaling (slope *b* ≈ 1.0), with axon size explaining the majority of the variance in vesicle count (*R^2^*= 0.69 and 0.56, respectively). In contrast, modulatory axons (acetylcholine and serotonin) displayed allometric scaling (*b* < 0.8) and weaker correlations, particularly for serotonergic axons (*b* = 0.48, *R^2^* = 0.22), suggesting that vesicle content in these boutons is less strictly constrained by axon calibre (Figure 4 - figure supplement 1).

Regions with high vesicle densities in glutamatergic and GABAergic axons indicated the presence of synapses, but this was not the case in the neuromodulatory axons. We identified synapses on the ultrastructural details available, including the presence of vesicles, and also any discernible membrane specialisation (Figure 5A-B). In acetylcholine and serotonin axons, the vesicle clustering was not as pronounced, and presynaptic specialisations, and postsynaptic densities, less distinct. To compare synapse sizes, the pre and postsynaptic density were segmented on each section, allowing us to extract the total area of this contact site. These were significantly smaller in the acetylcholine and serotonin axons (Figure 5C, synapse area µm^2^, median [Q1, Q3], glutamate: 0.20 [0.12, 0.28] µm^2^, *n* = 17, GABA: 0.11 [0.07, 0.18] µm^2^, *n* = 31 , acetylcholine: 0.03 [0.03, 0.04] µm^2^, *n* = 24, and serotonin: 0.03 [0.02, 0.04] µm^2^, *n* = 13; Mann–Whitney U test, glutamate to acetylcholine *U* = 666, *p* = 1.04 x 10^-6^; glutamate to serotonin *U* = 218, *p* = 4.52 x 10^-5^; GABA to acetylcholine *U* = 402, *p* = 3.80 x 10^-6^; GABA to serotonin *U* = 362, *p* = 2.32 x 10^-4^).

**Figure 5.**
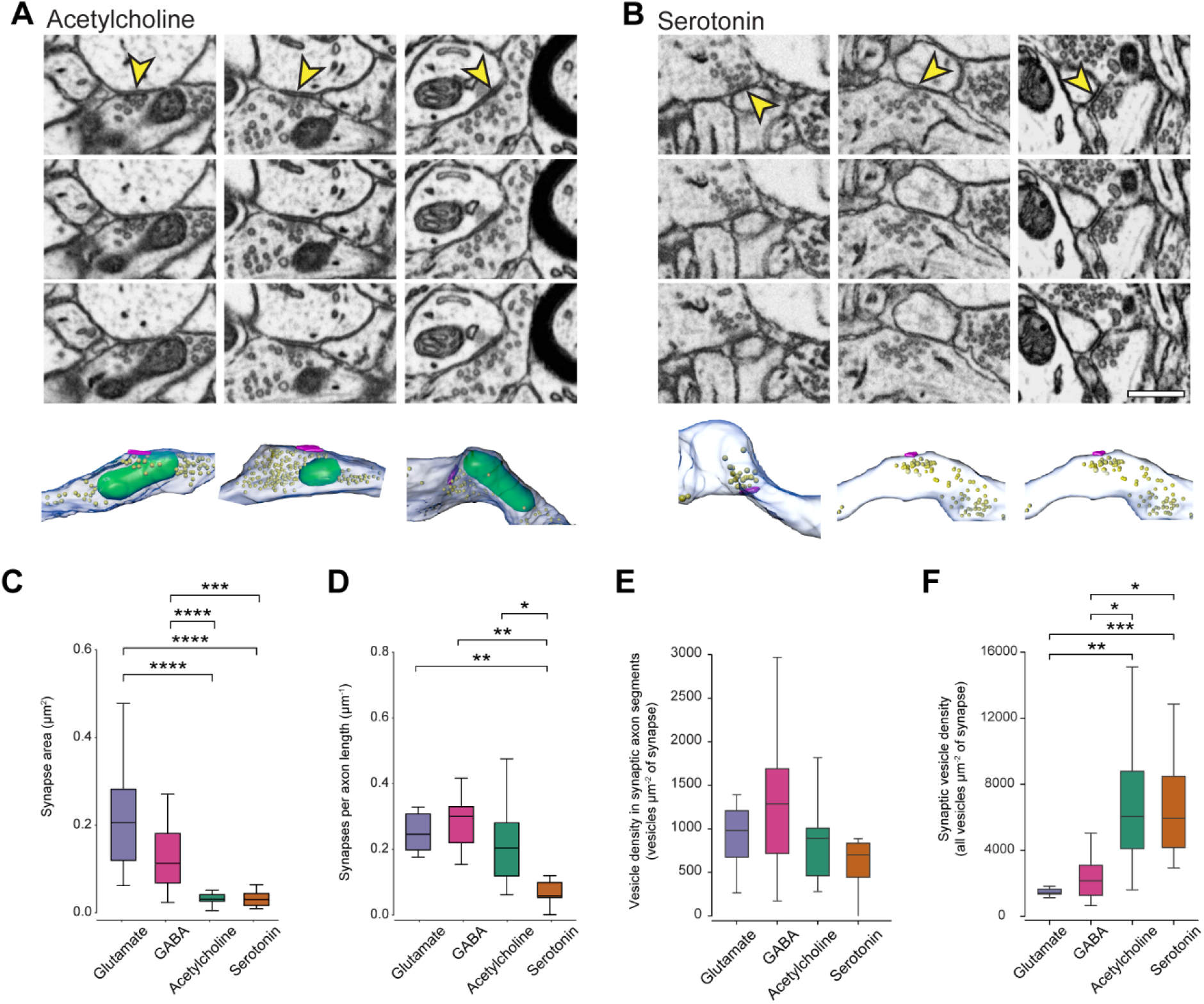
Ultrastructural identification and quantification of acetylcholine and serotonin synapses. **(A)** Serial electron micrographs of three representative cholinergic synapses. Yellow arrowheads indicate synaptic specialisations across consecutive sections. Corresponding 3D reconstructions of the same axonal segment with the synaptic contacts are shown below each example. **(B)** Serial electron micrographs of three representative serotonergic synapses, displayed as in **(A)**. Yellow arrowheads mark synaptic contacts across consecutive sections. Corresponding 3D reconstructions are shown below. Scale bar in lower right image in 0.5 µm. **(C)** Synapse area, measured as the area covered by the pre and postsynaptic density, of glutamate and GABA axons are larger than those of acetylcholine and serotonin. Median [Q1, Q3], n = number of synapses measured, glutamate: 0.20 [0.12, 0.28] µm^2^, n = 17, GABA: 0.11 [0.07, 0.18] µm^2^, n = 31, acetylcholine: 0.03 [0.03, 0.04] µm^2^, n = 24, and serotonin: 0.03 [0.02, 0.04] µm^2^, n = 13; Mann–Whitney U test, glutamate to acetylcholine *U* = 666, *p* = 1.04 x 10^-6^; glutamate to serotonin *U* = 218, *p* = 4.52 x 10^-5^; GABA to acetylcholine *U* = 402, *p* = 3.80 x 10^-6^; GABA to serotonin *U* = 362, *p* = 2.32 x 10^-4^. **(D)** The frequency of synapses along glutamate, GABA, and acetylcholine axons was similar, but significantly less in the serotonin type. Median [Q1, Q3], n = number of axons sampled, glutamate: (continued from D) 0.25 [0.20, 0.31] µm^-1^, n = 6, GABA: 0.30 [0.22, 0.33] µm^-1^, n = 9, and acetylcholine: 0.20 [0.12, 0.28] µm^-1^, n = 17, serotonin: 0.06 [0.05, 0.10] µm^-1^, n = 7; Mann–Whitney U test, glutamate to serotonin *U* = 42, *p* = 6.99 x 10^-3^; GABA to serotonin *U* = 63, *p* = 1.05 x 10^-3^; acetylcholine to serotonin *U* = 106, *p* = 1.14 x 10^-2^. **(E)** Vesicle density per unit synapse area, calculated by counting all vesicles in all 100 nm segments of axon where a synapse was present. Whisker box plot shows number of vesicles per µm2, median [Q1, Q3] n = number 100 nm segment where synapse(s) were present, glutamate 983.10 [675.73, 1209.09] n = 176, GABA 1286.50 [717.22, 1690.15] n = 141, acetylcholine 891.24 [461.82, 1008.08] n = 66, serotonin 701.80 [445.29, 836.14] n = 50; all comparisons were not significant (*p* > 0.05). Pairwise comparisons were performed using Mann-Whitney U statistics with Bonferroni correction for multiple comparisons (n = 6). All comparisons were not significant (*p* > 0.05). **(F)** Global vesicle density per unit synapse area, expressed as the ratio of total vesicle count to total synapse area calculated at the whole-axon level. Axons were included in the analysis only if they contained at least one synaptic region. Whisker box plot shows number of vesicles per µm^-2^ of synapse; median [Q1, Q3] n = number of axon segments, glutamate 1412.42 [1340.40, 1642.45] n = 11, GABA 2157.60 [1277.55, 3091.64] n = 10, acetylcholine 6851.82 [6042.52, 4099.58] n = 10, serotonin 5941.83 [4169.98, 8480.78] n = 7; Pairwise comparisons were performed using Mann-Whitney U statistics with Bonferroni correction for multiple comparisons (n = 6), Glutamate to acetylcholine *U* = 3.00 *p* = 1.72 x 10^-3^; glutamate to serotonin *U* = 0.00 p = 3.77 x 10^-4^; GABA to acetylcholine *U* = 14.00 *p* = 4.37 x 10^-2^; GABA to serotonin *U* = 5.00 *p* = 1.17 x 10^-2^.

However, the frequency that synapses appeared along the acetylcholine axons was similar to glutamate and GABA, but significantly less in the serotonin type (Figure 5D, synapses per axon length µm^-1^, median [Q1, Q3], glutamate: 0.25 [0.20, 0.31] µm^-1^, *n* = 6, GABA: 0.30 [0.22, 0.33] µm^-1^, *n* = 9, and acetylcholine: 0.20 [0.12, 0.28] µm^-1^, *n* = 17, serotonin: 0.06 [0.05, 0.10] µm^-1^, *n* = 7; Mann–Whitney U test, glutamate to serotonin *U* = 42, *p* = 6.99 x 10^-3^; GABA to serotonin *U* = 63, *p* = 1.05 x 10^-3^; acetylcholine to serotonin *U* = 106, *p* = 1.14 x 10^-2^). With fewer vesicles present per unit volume of the neuromodulatory axons (Figure 4C), along with smaller synapses, we questioned whether the number of vesicles present in the vicinity of putative release sites was in fact the same. To measure this, we used the centerline data (shown in Figure 3) and counted the numbers of vesicles in each 100 nm segment where a synapse was present, and then normalised this total to the total area of the synapse. Although this analysis only considers vesicles in the axon segments possessing a synapse, ignoring those that are synapse-free, this gives us a way of making an unbiased comparison of the abundance of vesicles at putative release sites. It also avoids having to make any estimates of where a varicosity may start or finish. This showed that despite the low numbers of vesicles in the neuromodulatory axons, per unit area of synapse, it is the same across all types (Figure 5E; number of vesicles per µm^2^ median [Q1, Q3] number of vesicles per 100 nm segment where synapse(s) were present, glutamate 983.10 [675.73, 1209.09] *n* = 176, GABA 1286.50 [717.22, 1690.15] *n* = 141, acetylcholine 891.24 [461.82, 1008.08] *n* = 66, serotonin 701.80 [445.29, 836.14] *n* = 50; all comparisons were not significant (*p* > 0.05)).

If, however, we ignore this spatial constraint, and make the assumption that all the vesicles in the axons could be involved in transmission at the synapses, then the comparison is very different (Figure 5F). In this case, the numbers of vesicles per unit area of synapse are far greater in the neuromodulatory axons (Figure 5F; number of vesicles per µm^2^, median [Q1, Q3] number of axon segments, glutamate 1412.42 [1340.40, 1642.45] *n* = 11, GABA 2157.60 [1277.55, 3091.64] *n* = 10, acetylcholine 6851.82 [6042.52, 4099.58] *n* = 10, serotonin 5941.83 [4169.98, 8480.78] *n* = 7; Mann-Whitney U statistics with Bonferroni correction for multiple comparisons (*n* = 6), Glutamate to acetylcholine *U* = 3.00 *p* = 1.72 x 10^-3^; glutamate to serotonin *U* = 0.00 *p* = 3.77 x 10^-4^; GABA to acetylcholine *U* = 14.00 *p* = 4.37 x 10^-2^; GABA to serotonin *U* = 5.00 *p* = 1.17 x 10^-2^).

This large discrepancy between the two types of axon (Figure 5F) highlights the large proportion of vesicles in neuromodulatory axons that are very far from the sites of release. Or at least far from the structural specialisations that we have identified as synaptic release sites. This led us to consider what proportion of the vesicles, in the different axons, that are close to the membrane, were in fact associated with a synapse? If vesicles of glutamate and GABA axons were in the regions where there were synapses, what proportion of their vesicles were in fact close to these sites of release. And how different is this in the neuromodulatory axons? In all the reconstructed axons, vesicles had also been plotted (examples shown in Figure 6A) so we could calculate the closest distance to the plasma membrane, but also to the nearest synapse. We only considered distances of less than 180 nm, and these were grouped into 30 nm distance bins (0 - 30; 30 - 60; 60 - 90; 90 - 120; 120 - 150; 150 - 180 nm; Figure 6B, C). This analysis shows the extent to which vesicles in the neuromodulatory axons are predominantly located far from any synaptic specialisations (Figure 6B, C). If we consider only the vesicles that are within 180 nm of the plasmalemma or synapse, then in glutamate and GABA axons, 43.8 and 39.2 % are closer to synapses. However, in the neuromodulatory axons this is only 7.9 % of vesicles in acetylcholine, and 10.7% for serotonin axons (Figure 6B, C). This shows the vast majority of vesicles being far from synapse, but what about those that are at the sites of release? Are those that are close to the synapse arranged in a similar way to those in the fast transmitters? The isotropic resolution of the FIBSEM dataset (5 nm voxel size) enabled vesicles to be visualized across multiple consecutive sections, and allowed us to plot the position of all vesicles in 3D (Figure 6D). Here, we measured the distances of vesicles to the presynaptic membrane, and when normalised to the area of the synapse, the organisation looks very similar (Figure 6E, F). We considered those within 60 nm of the membrane to be docked (Figure 6E; *N*, Mean ± SEM, glutamate 32, 132.33 ± 14.65; GABA 31, 193.51 ± 18.74; acetylcholine 24, 147.78 ± 15.49; serotonin, 13, 157.99 ± 18.17), and here again, the neuromodulatory axons are similar to both the glutamate and GABA. However, GABA docked vesicles were more numerous to glutamate (*t-test* Bonferroni Adjusted, glutamate vs. GABA *p* = 0.04).

**Figure 6.**
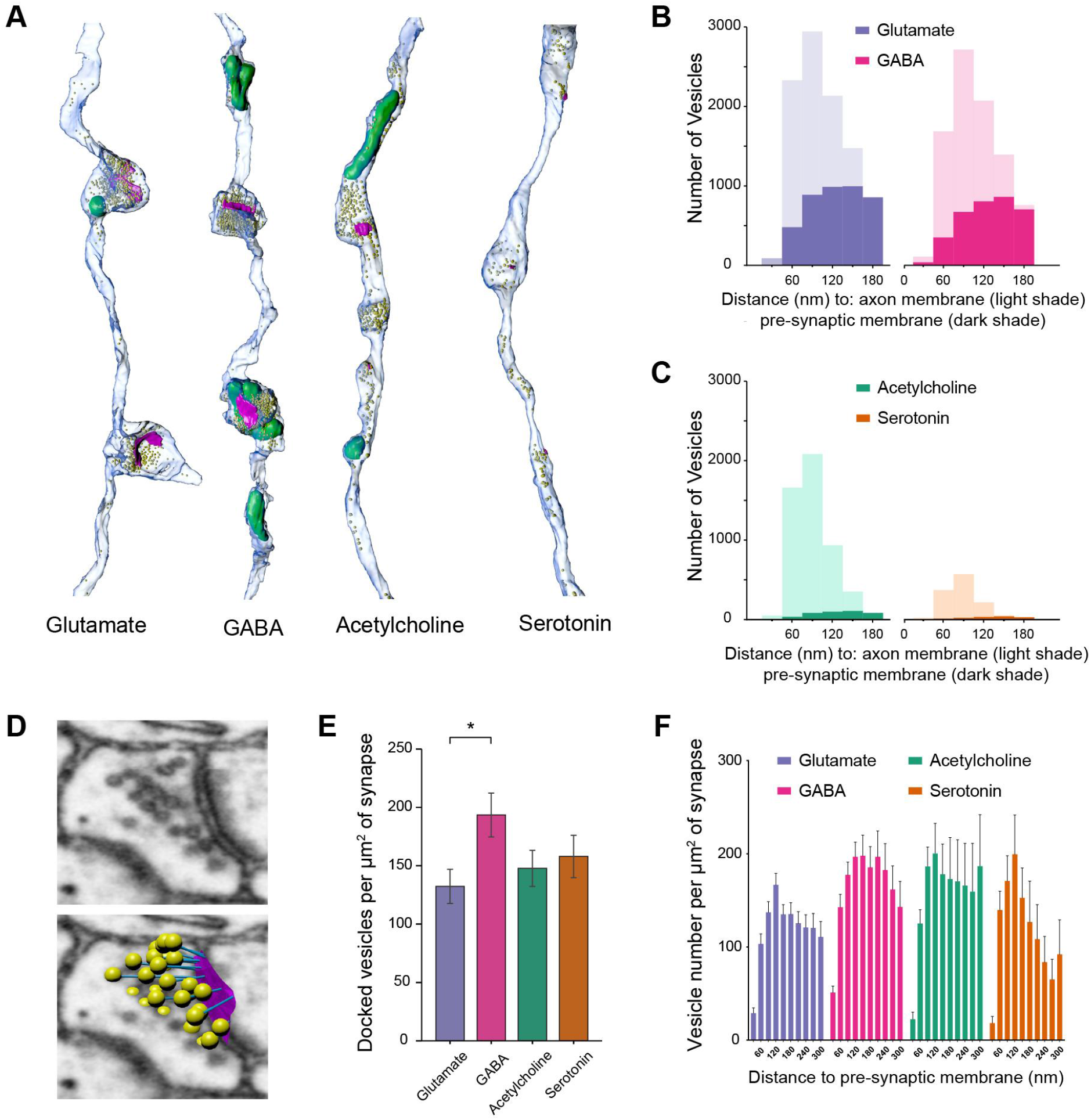
Vesicle proximity to the axonal membrane and the synapse. **(A)** Three-dimensional reconstructions of representative glutamate, GABA, acetylcholine, and serotonin axonal segments. Axonal membranes are rendered in translucent blue, vesicles as yellow spheres, synapses in magenta, and mitochondria in green. **(B-C)** Histograms of vesicle distances relative to the synapse (solid colour) and plasma membrane (shaded colour) for glutamate, GABA, acetylcholine and serotonin vesicles in 30 nm distance bins. The percentage of all vesicles that were within 180 nm of the plasma membrane, or synapse, was for glutamate vesicles, 89.2 % (9790 / 10978), GABA, 94.9 % (8732 / 9206), acetylcholine, 99.8 % (5136 / 5148) and serotonin, 99.9 % (1230 / 1231). The percentage of these vesicles that were closer to synapses than to the plasma membrane was for glutamate, 43.8 % (4,291 / 9,790), GABA, 39.2 % (3,426 / 8,732), acetylcholine, 7.9% (408 / 5136) and serotonin, 10.7 % (131 / 1230). Glutamate: (continued from B-C) 10978, 4291 (39.1%), 9790 (89.2%); GABA: 9206, 3426 (37.2%), 8732 (94.9%); acetylcholine: 5148, 408 (7.9%), 5136 (99.8%); serotonin: 1231, 131 (19.6%), 1230 (99.9%). **(D)** Electron micrograph of a presynaptic varicosity (top) and overlaying vesicle plots (bottom), illustrating vesicle positions (yellow) and the active zone (magenta). **(E)** Quantification of docked vesicle density (within 60 nm of the synapse) calculated as vesicles per unit surface area of synapse, across neurotransmitter classes. Bars represent mean ± s.e.m., and n = number of synapses sampled, Glutamate 132.33 ± 14.65, n = 32; GABA 183.51 ±18.74, n = 31; acetylcholine 147.78 ± 15.49, n = 24; serotonin 157.99 ± 18.17, n = 13. Student’s t-test with Bonferroni correction for multiple comparisons (n = 3). Glutamate to GABA *p* = 0.04. All other comparisons were not significant (*p* > 0.05). **(F)** Vesicle densities at 30 nm intervals from the presynaptic membrane, calculated as numbers of vesicles per unit surface area of synapse show similar profiles across the four neurotransmitter types.

The quality of the FIBSEM imaging (Figure 7A) not only allowed for the positions of vesicles to be plotted in 3D, but we could also estimate their size and shape by measuring their cross-sectional area in the single image plane in which each vesicle appeared largest. Using this measurement, the vesicle cross-sectional area differed significantly across transmitter classes (Figure 7B; median [Q1, Q3] *n*, glutamate 1632.14 [1404.21, 1845.51] µm^2^ *n* = 500, GABA 1731.19 [1518.72, 1965.28] µm^2^ *n* = 600, acetylcholine 2344.65 [2070.89, 2603.96] µm^2^ *n* = 600, serotonin, 2121.58 [1849.50, 2447.12] µm^2^ *n* = 600; Mann-Whitney U test with Bonferroni correction for multiple comparisons, Glutamate to GABA *U* = 121551, *p* = 3.43 x 10^-7^; glutamate to acetylcholine *U* = 24279 *p* = 4.5 x 10^-126^; glutamate to serotonin *U* = 51317.50 *p* = 3.80 x 10^-78^; GABA to acetylcholine *U* = 46135.50 *p* = 2.15 x 10^-109^; GABA to serotonin *U* = 85389 *p* = 3.42 x 10^-55^; acetylcholine to serotonin *U* = 225342 *p* = 2.54 x 10^-13^). Vesicles associated with acetylcholine and serotonin exhibited larger average areas and broader distributions than glutamatergic and GABAergic vesicles, indicating increased vesicle size and heterogeneity in neuromodulatory axons. In contrast, glutamate and GABA vesicles clustered at smaller areas with narrower distributions, consistent with more uniform vesicle populations.

**Figure 7.**
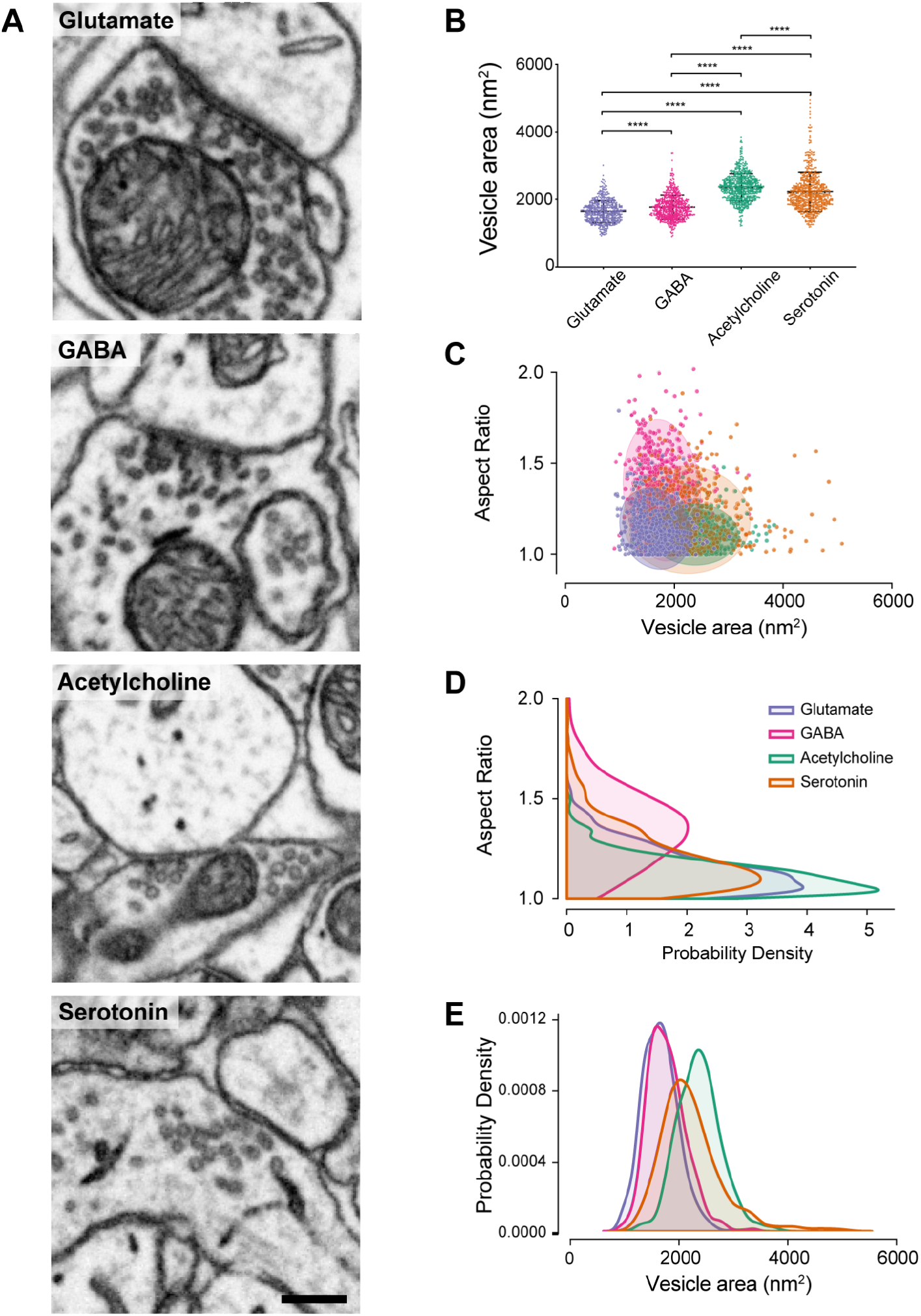
Shape of synaptic vesicles across neurotransmitter classes. **(A)** Representative electron micrographs of axonal varicosities of glutamate, GABA, acetylcholine and serotonin axons, showing synapses, and synaptic vesicles. Scale bar bottom right = 200 nm. **(B)** Distribution of vesicle cross-sectional area (nm^2^) across neurotransmitter classes. Swarm plot detailing the distribution of vesicle cross-sectional areas. Visually, vesicles belonging to the glutamate and GABA populations exhibit tighter clustering at lower values whereas acetylcholine and serotonin population show higher central tendencies and much larger variance, particularly in the serotonin group (see pronounced upper tail of high-value outliers). Each point represents a single vesicle. Overlaid horizontal black lines indicate the median [Q1, Q3] n = number of vesicles, glutamate 1632.14 [1404.21, 1845.51] µm^2^ n = 500, GABA 1731.19 [1518.72, 1965.28] µm^2^ n = 600, (continued from B) acetylcholine 2344.65 [2070.89, 2603.96] µm^2^ n = 600, serotonin, 2121.58 [1849.50, 2447.12] µm^2^ n = 600. Mann-Whitney U test with Bonferroni correction for multiple comparisons. Glutamate to GABA *U* = 121551, *p* = 3.43 x 10^-7^; glutamate to acetylcholine *U* = 24279 *p* = 4.5 x 10^-126^; glutamate to serotonin *U* = 51317.50 *p* = 3.80 x 10^-78^; GABA to acetylcholine *U* = 46135.50 *p* = 2.15 x 10^-109^; GABA to serotonin *U* = 85389 p = 3.42 x 10^-55^; acetylcholine to serotonin *U* = 225342 *p* = 2.54 x 10^-13^. **(C)** Morphological distribution of synaptic vesicles. Scatterplot illustrating the relationship between vesicle area and aspect ratio for four neurotransmitter populations. Each point represents an individual vesicle (all neurotransmitters except Glutamate, n = 600; Glutamate, n = 500). Confidence ellipses (2σ) represents the covariance structure for each group: Glutamate (purple), GABA (pink), acetylcholine (green) and serotonin (orange). The centroids indicate the population mean for each neurotransmitter subtype. The vertical elongation of the GABA ellipse reflects high variance in their aspect ratio (0.038), while the horizontal spread of the serotonin ellipse denotes high variance in vesicle area (3.4 x 105 nm^2^). The near-zero correlation coefficients calculated using Pearson’s (r = -0.017 to -0.153) and lack of diagonal orientation in all groups suggests that vesicle size and shape are regulated by independent mechanisms. **(D)** Kernel density estimates (x axis) of vesicle aspect ratio distributions (y axis) shown in **(D)** for each neurotransmitter class. GABA vesicles display a broader distribution with a pronounced shift toward higher aspect ratios, indicating a greater prevalence of elongated vesicles. In contrast, glutamate, acetylcholine, and serotonin vesicles have a predominantly more circular profile with aspect ratios of 1.0–1.2. **(E)** Kernel density estimates (y axis) of vesicle cross-sectional area distributions (x axis) shown in **(D)** for each neurotransmitter class. Glutamate and GABA show similar distributions with smaller profiles in comparison to acetylcholine vesicle areas that are shifted towards larger sizes. The serotonergic vesicles show an intermediate distribution reflecting increased variability of vesicle area.

Vesicle shape was also assessed using their aspect ratio, measured from the same maximal cross-sectional plane, providing a two-dimensional estimate of deviation from circularity (Figure 7C, D, E). While all four vesicle populations share a similar central tendency (represented by the centroid of each ellipse, and demonstrated by their overlapping means), they differ in their variability. Glutamate and acetylcholine vesicles form tight, compact populations with low variances in both dimensions (*e.g.,* acetylcholine aspect ratio variance, 0.007). GABA vesicles, on the other hand, show a substantial variation in shape, with a high aspect ratio variance (0.038), which is two to three times greater than that of the other groups. Despite this, its area remains comparable to the other groups, resulting in a pleomorphic appearance. In contrast, serotonin vesicles vary widely in size, with the largest area variance (3.4 x 10^5^ nm^2^) but maintain relatively consistent shapes, suggesting a looser regulation of vesicle volume. Across all four types of vesicle the ellipses are not orientated diagonally. This aligns with Pearson correlations (r) being close to zero (-0.017 to -0.135), indicating that the mechanisms regulating vesicle size and shape are independent.

Plotting vesicle positions near synapses allows comparison of their spatial organisation between the axon types, but this depends on the reliable identification and segmentation of synaptic contacts. While synapses are easily identified in glutamate and GABA axons, they appear smaller and less distinct in the neuromodulatory type. Together with the wide distribution of vesicles along these axons, this prompted us to test whether vesicle clusters assigned to synapses represent the dominant structural feature of axon wiring. Specifically, are vesicles preferentially concentrated at the contact sites that we assigned as synapses rather than distributed elsewhere, and conversely, other elements in the neuropil? To address this, we looked at the correlation between vesicle numbers in the axon, at 100 nm intervals, and the amount of the different cellular elements that were touching this 100 nm segment (Figure 8A, B). The different touching elements measured were: dendrites, dendritic spines, astrocytic processes, other axons, and identifiable synapses.

**Figure 8.**
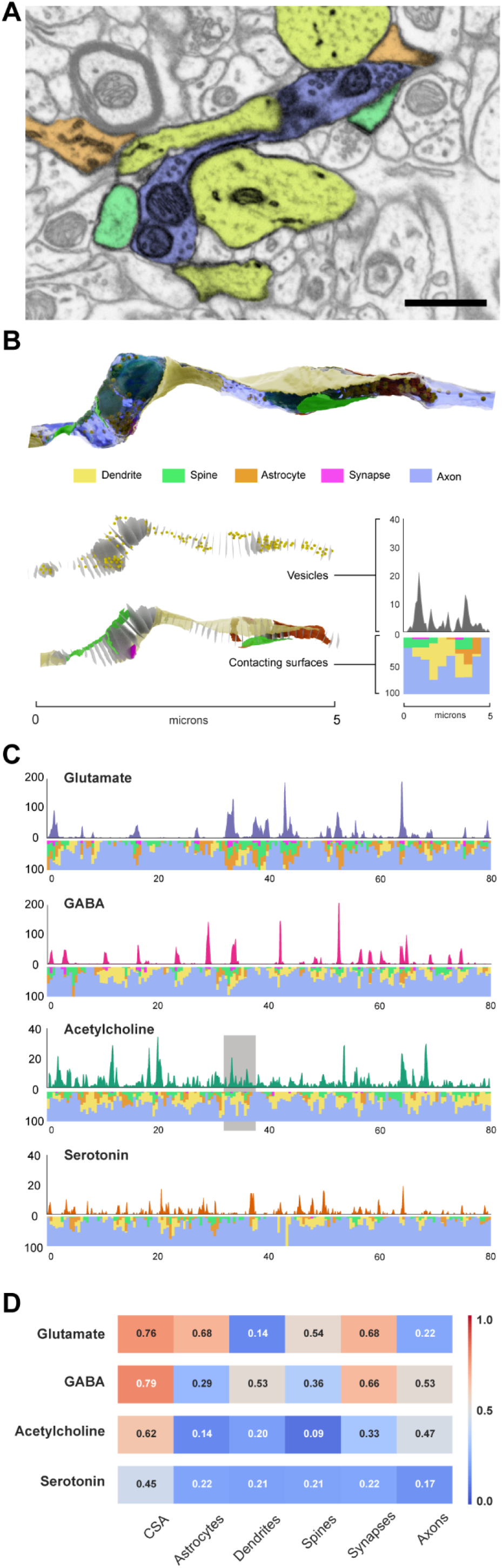
Relationship between vesicle number and membranes in apposition with individual axons. **(A)** Representative FIBSEM image from a serial image stack through a cholinergic axon segment. Cellular elements in apposition with the reconstructed axon are pseudo-coloured to illustrate the surrounding neuropil. The axon of interest is coloured blue and is contacted by dendrites (yellow), dendritic spines (green), and astrocytic processes (orange). Other axons contacting the blue axon have been left uncoloured. **(B)** Three-dimensional reconstruction of the axon shown in **(A)**, generated from the serial FIBSEM dataset. Upper panel: reconstructed axon showing the membranes of surrounding elements in apposition using the same colour scheme as in **(A)**. Middle panel: distribution of synaptic vesicles (yellow balls) along the axon length, and cross-sections of axon positioned at 100 nm intervals. Right hand plot shows the vesicle number along the length of the axon shown. Lower panel: membranes in apposition with the axon, and cross-sectional areas. Right hand plot shows the percentage coverage of each 500 nm segment of dendrites, dendritic spine, astrocyte, synapse, and other axons using the same colour scheme as above. **(C)** Linear analysis of 80 µm length samples for each type of axon, as shown in **(B)**, right hand graphs. The upper trace shows vesicle number measured at 100 nm intervals along the axon length, while the stacked coloured bars below indicate the proportion of membrane contact contributed by surrounding cellular elements at 500 nm intervals. Colour scheme is as above: dendrites (yellow), astrocytes (orange), dendrites (yellow), dendritic spines (green), synapses (magenta), and axons (blue). **(D)** Heat map showing correlation coefficients between vesicle number and area of contact of different cellular elements at 100 nm intervals along axon length. Correlations are shown for axon cross-sectional area (CSA) and for membrane contact by astrocytes, dendrites, dendritic spines, synapses, and axons. Warmer colours indicate stronger positive correlations, whereas cooler colours indicate weaker correlations, as shown by the scale bar. Values within each cell indicate the corresponding coefficient (r). The scale bar in **(A)** is 0.5 µm.

In glutamate axons, the number of vesicles exhibited a strong positive correlation with the amount of synaptic membrane (r = 0.68; Fig. 8D), showing, as expected, that vesicles are preferentially concentrated at sites associated with postsynaptic specialisations. There were also high correlations with dendritic spines (r = 0.54), and astrocytic membranes (r = 0.68), the site of glutamate reuptake. Similarly, for GABA axons, the correlation was high for the number of vesicles and their synaptic contacts (r = 0.66), and membranes of the dendritic shaft (r = 0.52), but not spines (r = 0.36), showing their preference of synaptic partner. In contrast, however, this coupling was substantially weaker in acetylcholine (r = 0.33) and serotonergic (r = 0.22) axons, indicating that vesicle accumulation in these axons is not tightly linked to synaptic membrane specialisations, or indeed any of the other surrounding cellular elements - a clear indication that vesicles in neuromodulatory axons were not accumulating at other sites along the axon. In this analysis the strongest correlation in neuromodulatory axons was with cross-sectional area (acetylcholine r = 0.62; serotonin r = 0.45), consistent with the observation that vesicles are distributed more diffusely along axonal swellings rather than concentrated at discrete release sites. To understand if the vesicles are randomly distributed, we performed simulations using a simplified axonal geometry embedded in neuropil. These synthetic axons were cylinders placed in the serial image volumes, and again, the cellular elements in juxtaposition with them were reconstructed, in the same way as the real axons. Vesicle distributions from reconstructed acetylcholine and serotonin axons were placed within these cylinders and their position randomised repeatedly in simulation while maintaining the original vesicle counts for each axon (Figure 8 - figure supplement 1). The resulting correlation matrix demonstrates an absence of strong linear relationships, with coefficients between -0.08 and 0.22, consistent with a random spatial distribution.

These findings demonstrate that, unlike classical excitatory and inhibitory synapses, vesicle organisation in cholinergic and serotonergic axons is weakly coupled to synaptic membrane specialisations, supporting a model in which neuromodulatory transmission is supported by a more spatially diffuse and structurally distinct axonal architecture.

## Discussion

A central question in neuromodulation is how the anatomical organisation of long range modulatory axons supports their mode of signalling within cortical circuits. Unlike the fast transmitters, glutamate and GABA, acetylcholine and serotonin are widely thought to act, at least in part, via volume, or non-synaptic, transmission; however, the structural basis for this mode of communication has largely been inferred from ultrastructural studies that rely on immunocytochemical labelling to identify axons of interest, and by only targeting varicosities with vesicles or synaptic profiles (*e.g.,* serotonin: Beaudet & Descarries, 1976; Descarries et al., 1975; Kosofsky & Molliver, 1987; Papadopoulos et al., 1987; Séguéla et al., 1989; Smiley & Goldman-Rakic, 1996; and acetylcholine: Mrzijak et al., 1995; Smiley et al., 1997; Turrini et al., 2001; Umbriaco et al., 1994). While these approaches have provided valuable information, the use of electron dense reaction products, such as diaminobenzidine, can obscure fine subcellular features, including vesicle populations and membrane specialisations. This limits the precision with which axonal architecture can be resolved. Additionally, by focusing the analysis on sites of axon swelling, varicosities, or synaptic sites, these studies necessarily sample only a subset of the axon, biasing interpretation toward regions enriched in vesicles or structural specialisations. As a result, these analyses have provided no information about the distribution of the actual structures involved in transmission (*i.e.* the vesicles), and therefore say nothing about how the population is distributed along the extent of the axon, or how these distributions relate to the surrounding neuropil and potential sites of release. The analyses presented here were centered around extended segments of genetically identified axons, first selected with light microscopy, and not electron microscopy on the basis of varicosities or synaptic specialisations. In this way we could capture whole pieces of axon and then plot the position of all vesicles contained in a synapse and varicosity-agnostic manner. This approach enables an unbiased assessment of the spatial relationship between vesicles and the axonal plasmalemma, including identifiable release sites, and allows us to determine the extent to which vesicles may or may not be preferentially associated with synaptic structures.

This is the first time that such an approach has been used, and even a cursory look at the light microscopy images and their corresponding 3D reconstructions (Figure 1 - figure supplement 1) shows how difficult it would be to predict synaptic connectivity based on the presence of boutons or varicosities, in the way that is possible with glutamate (Gala et al., 2017) and GABA axons (Wu et al., 2012). Our analyses of this geometry alone suggests that the neuromodulatory axons are organised according to different principles than canonical excitatory and inhibitory axons. As well as their lack of clear varicosities, the vesicle abundance, vesicle morphology, and their relationship to synaptic specialisations are very different, and support the idea, drawn from previous EM analyses, that neuromodulatory transmission occurs through a distinct axonal architecture rather than any subtle differences of classical synaptic transmission.

The previous analyses, that found smaller synaptic connections, and inferred lower concentrations of these release sites by targeting the axon swellings (Beaudet & Descarries, 1976; Descarries et al., 1975; Umbriaco et al., 1995), came to the conclusion that rather than there being a simple dichotomy between synaptic and nonsynaptic modes of transmission, neuromodulatory signalling might operate along a spectrum of release architectures. Although our current data could support this idea, it does show that the specialisations, we identify as synapses, have a remarkably low proportion of the entire vesicle population associated with them. This is only 10% in the case of acetylcholine and 7% in the case of serotonin. Therefore, so few vesicles associated with indistinct synaptic structures, suggests limited importance of point-to-point synaptic transmission, and a far more dominant release mechanism elsewhere, ie. non synaptic, volume transmission. In addition to the possibility of very limited synaptic transmission we must also bear in mind the structures identified in these and previous analyses only ever show weak hallmarks that typically define synapses (Gray, 1959a, 1959b). In all instances, membrane specialisations were indistinct and small, and vesicle clustering were small, and faintly defined, making unambiguous synapse identification difficult. As none of the connections were as defined as those of the glutamate and GABA axons, they could represent the lower end of a continuum of synaptic specialisations, however, our analysis of the arrangement of vesicles at these synapses may not support this. Ultrastructural studies of fast transmitting axons have well-established the correlation between synapse size and vesicle number. Larger presynaptic boutons with bigger active zones generally contain more docked and total synaptic vesicles, whereas smaller boutons contain fewer in glutamate axons at least (*e.g.,* Santuy et al., 2018; Schikorski & Stevens, 1997). As the lack of discernible varicosities precludes any analysis based on this structure, we could measure the area of synapse, and number of vesicles, at 100 nm intervals along the axons’ length. Like previous analyses (Eyre et al., 2007; Nusser et al., 1997; Schikorski & Stevens, 1997) this showed a correlation in the glutamate and GABA synapses, but for acetylcholine and serotonin we show this as being weak. So not only do we have smaller and less distinct synapses, but the amount of vesicles gathered at these sites doesn’t scale in the same proportion as we see with the fast transmitters. Therefore, we could consider the possibility that the structures we designate as synapses may not be release sites at all, but junctional adhesion specialisations. During development, similar junctions are precursors of synaptic connections, and a number of EM studies in adult brain describe junctional complexes with symmetric cytoplasmic densities consistent with adherens junctions positioned at synaptic margins or adjacent to synapses (Spacek & Harris, 1998; Uchida et al., 1996). Synaptic junctions have been proposed to include a distinct catenin-based adherens junction subdomain bordering the transmitter release zone.

If, therefore, the neuromodulatory axons have such an inconsequential synaptic release, then the location of vesicles, rather than the synapses themselves, would be a better indication of their function. Correlating the numbers of vesicles with the identity of the cellular elements juxtaposed to the axons shows expectedly high correlations in the glutamate and GABA axons. In the case of glutamate; spines, the synapses themselves, as well as astrocytes; and for GABA, dendrites and also their synapses. For neuromodulatory axons there is little indication of any relationship with any particular cellular element, apart from, however, acetylcholine axons which show a moderate correlation with neighbouring axons, and the same correlation as GABA axons. While this finding raises the possibility that neighbouring axons may represent sites of presynaptic modulation, consistent with established axo-axonic actions of acetylcholine and GABA (Bowery et al., 1980; Kawai et al., 2007; Lambe et al., 2003). This association of vesicle-rich regions with other axons raises the possibility that these sites may contribute to local presynaptic modulation rather than signalling that is directed towards a specific postsynaptic target.

The correlations with neighbouring axons, though weak, do not preclude the idea that vesicle localisation is organised with respect to surrounding cellular elements. This was addressed by carrying out the same analysis on artificial axons which were placed inside the same FIBSEM image volumes, with membranes of apposed cellular elements reconstructed on their surface. Here, the correlations are substantially lower (Figure 8 - figure supplement 1). This indicates that vesicle localisation in neuromodulatory axons is not simply a consequence of the local geometry of the neuropil, but exhibits a degree of biological organisation. While this supports the existence of an underlying organisational principle, the absence of any conventional synaptic structures suggests that the organisation is unlikely to reflect classical target-specific synaptic connectivity. Instead, it points towards a broader neuromodulatory signalling with axons regulating network state rather than transmitting discrete signals to individual postsynaptic partners.

The differences in the structural organisation between neuromodulatory and fast transmitting axons were also seen at the level of the vesicles themselves. Vesicles associated with acetylcholine and serotonin were larger and more heterogeneous in shape than those associated with glutamate and GABA, which were smaller and in the case of glutamate, more uniformly spherical. Differences in vesicle size in monoaminergic axons have been noted qualitatively in previous EM studies (Descarries et al., 2004; Nirenberg et al., 1995). Our quantitative analysis demonstrates that vesicle morphology itself is a distinguishing feature of neuromodulatory axons. Although the shape of vesicles is understood to be influenced by the chemical fixation process used to prepare the brain tissue for EM (Korogod et al., 2015), this could also be a reflection of distinct molecular compositions of the different types of vesicles, and another indication of neuromodulatory transmission using very different release machinery to that used for fast synaptic transmission.

These current findings show that the structure and organisation of neuromodulatory axons differ in important ways from that of classical excitatory and inhibitory axons. Vesicle organization in cholinergic and serotonergic axons is diffuse, weakly coupled to synaptic membrane specializations, and associated with distinct vesicle morphologies. While these observations do not preclude the formation of conventional point-to-point synaptic connections in neuromodulatory axons, morphological evidence suggests that fast connections may represent only a tiny subset of their signaling repertoire. Their features support models of neuromodulatory transmission that are biased towards a spatially distributed release, and volume transmission plays the dominant role in their signalling. By providing quantitative, three-dimensional evidence for these differences, this work offers a structural foundation for interpreting how neuromodulatory systems exert widespread influence in cortical circuits.

These findings also have important implications for large-scale connectomic efforts that rely on ultrastructural criteria to define synaptic contacts. These criteria are optimised for glutamate and GABA connections, where vesicle clustering, active zones, and postsynaptic densities define synapses. However, in neuromodulatory axons, the weak expression of these features, together with the poor correlation between vesicle distribution and specific cellular partners, suggests that conventional synapse definitions may miss, or at least underestimate sites of communication. Wiring diagrams derived from EM reconstructions capture glutamatergic and GABAergic connectivity but could fail to correctly represent the neuromodulatory signalling component. Our vesicle-centric, synapse-agnostic approach highlights the need for alternative annotation strategies that incorporate vesicle distributions, axon–environment relationships, and molecular identity, in order to more accurately reflect the organisational principles of neuromodulatory systems in connectomic datasets.

## Methods

### Animal experimentation

All experimentation on animals was undertaken according to Swiss federal regulations and strictly in accordance with local authorities of animal experimentation under licences issued to Anthony Holtmaat (Licence: GE/74/18), University of Geneva, and Carl Petersen (Licence: VD1628), EPFL.

### Labelling of Cholinergic axons

Cholinergic axons were labelled using AAV-FLEX-GCaMP6s-mRuby (1.5 µL, 1.83 × 10^13^ GC, Addgene 68717-AAV1) (Rose et al., 2016) with injections targeting the center of the basal forebrain using the following coordinates from bregma: AP, −0.5 mm; ML,1.8 mm, DV, −4.5 mm. Injections were made in 10-12 weeks old adult mice positive for ChAT-Cre genotyping (JAX: 031661) (Rossi et al., 2011).

### Labelling of Serotonergic axons

Serotonergic axons were labelled using AAV-CAG-Flex-tdTomato-WPRE-bGH virus (210 nL, 1.83 × 10^13^ GC, Penn Vector Core) with injections targeting the median and dorsal raphe nuclei (DRN and MRN) using the following coordinates from bregma: AP,−4.50 and −4.80 mm; DV, −4.50, -3.50 and -3.00 mm. Injections were made in 10-12 weeks old adult mice, positive for both 5HT3aR-BAC^EGFP^ and SERT^cre^ genotyping.

### Tissue fixation and 2-photon imaging

Mice were deeply anaesthetised with an intraperitoneal injection of ketamine (0.13 mg g^−1^ body weight) and xylazine (0.01 mg g^−1^) and transcardially perfused with phosphate buffered saline (0.01M, pH 7.4), immediately followed by a buffered mix of 1.2% glutaraldehyde and 2% paraformaldehyde in 0.1 M phosphate buffer. After perfusing with a minimum of 30 mL per animal they were left for 2 hours before the brain was removed ready for sectioning. Vibratome sections were cut at 80 micron thickness, and tangential to the somatosensory cortex (VT1200, Leica Microsystems).

Two photon imaging was performed on the fixed sections using a custom-built 2PLSM, equipped with a femtosecond-pulsed titanium-sapphire laser (Chameleon Ultra II, Coherent) and a 20× water immersion objective (0.8 NA, Zeiss). Both low-resolution image stacks (512×512 pixels; 3.2 µs pixel dwell time; 0.3 µm/pixel; 3 µm steps) of the region of interest and high-resolution image stacks (512×512 pixels; 3.2 µs pixel dwell time; 0.09 and 0.05 µm/pixel; 1 µm steps) of selected axons in cortical layer 2 were collected and then fiducial marks made by burning lines into the fixed tissue, using the laser (Maco et al., 2014). GFP-expressing axons were imaged at λ= 910 nm, tdTomato-expressing neurons were imaged at λ = 1040 nm.

### Preparing brain slices for electron microscopy

To prepare the selected sections for electron microscopy, they were first washed thoroughly with cacodylate buffer (0.1 M, pH 7.4), then postfixed for 40 minutes in 1.0% osmium tetroxide with 1.5% potassium ferrocyanide, followed by 40 minutes in 1.0% osmium tetroxide alone. They were finally stained for 40 minutes in 1% uranyl acetate in water before being dehydrated through increasing concentrations of ethanol and then embedded in Durcupan ACM (Fluka, Switzerland) resin between two glass slides coated in mold releasing agent.

Once the resin had hardened, it was released from between the two glass slides using a razor blade, and the regions of interest in the sections located. These were outlined by lightly scratching a mark on the resin surface and were then cut away from the rest of the section and glued to a blank resin block. This was trimmed down, using glass knives mounted in an ultramicrotome. The orientation of the block face was such that it was parallel to the plane of the brain section which was parallel to the imaging plane of the light microscope. After trimming the region of interest was positioned close to the edge of the block so that the focussed ion beam could be directed parallel to the block face, and the region of interest, and did not mill any unwanted part surrounding it. This ensured that the milling was consistent with consistent amounts of resin being removed each time before the imaging.

### Milling and Imaging

Blocks were milled and imaged inside a Zeiss NVision 40 FIBSEM (Carl Zeiss SMT, Germany). To achieve consistent milling and imaging through the face, a protective layer (∼ 0.5 µm) of carbon was deposited onto the surface of the block, above the region of interest. An acceleration voltage of 2 kV, with a current of between 340–400 pA and dwell time of 10 µs/pixel were used. Each image was collected at 5 nm per pixel with each image. The imaging and milling cycle was maintained at below 2 minutes, with a total image size of 2048×1536 pixels (∼3 MB/image). This corresponds to a field size of 12.45 µm×9.34 µm. The milling depth after each image was approximately 7 nm , and for each stack was determined precisely using the cylindrical diameters method (Fiala & Harris, 2001). This method calculates the section thickness by averaging the ratios of the diameter of mitochondria sectioned longitudinally within the imaging plane to the number of sections they span. This was done across multiple regions containing mitochondria.

### Image Processing and 3D Reconstruction

The final image stack was aligned in the Fiji software package (http://fiji.sc/wiki/index.php/Fiji) and the axons of interest identified using the 2 photon image stacks to compare features. Once found, segmentation of the axons, their synapses and mitochondria were drawn, as well as the position of every vesicle plotted, using the TrakEM2 plugin in the same software (Cardona et al., 2012). Additionally, every membrane in apposition with the drawn axon was drawn. These surrounding membranes belonged to other structures in the neuropil: axons, dendrites, astrocytes, or cell bodies. No oligodendrocytes, or microglial membranes were identified.

### Analysis of the 3D models

Once the axons, and associated structures, had been segmented, 3D models were exported from the TrakEM2 software and imported into the Blender software (www.blender.org). Morphometric measurements were obtained from the mesh-based models using the NeuroMorph tools in Blender (Jorstad et al., 2018). The software computes geometric parameters such as surface area, volume, linear distances, and cross-sectional area, directly from three-dimensional mesh reconstructions, providing quantitative characterization of the analyzed structures. Centrelines were calculated using the VMTK software (Antiga et al., 2008), Vascular Modeling Toolkit; RRID:SCR_001893).

## Statistics

Mann-Whitney U with Bonferroni’s correction for multiple comparison. Significance is denoted by brackets: **** (*p* < 0.0001), *** (*p* < 0.001), ** (*p* < 0.01), and * (*p* < 0.05).

## Supplementary Figures

**Figure 1—figure supplement 1.**
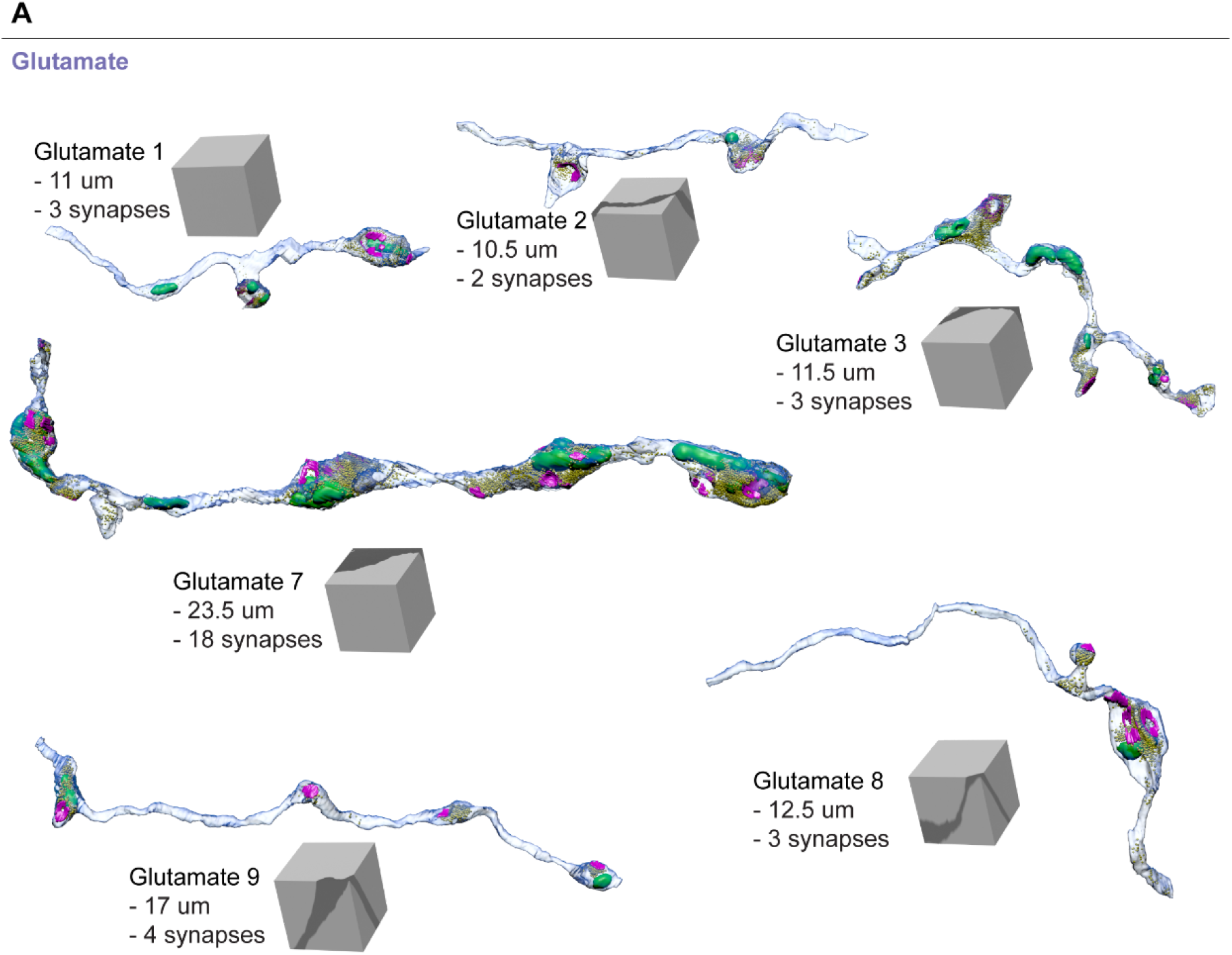
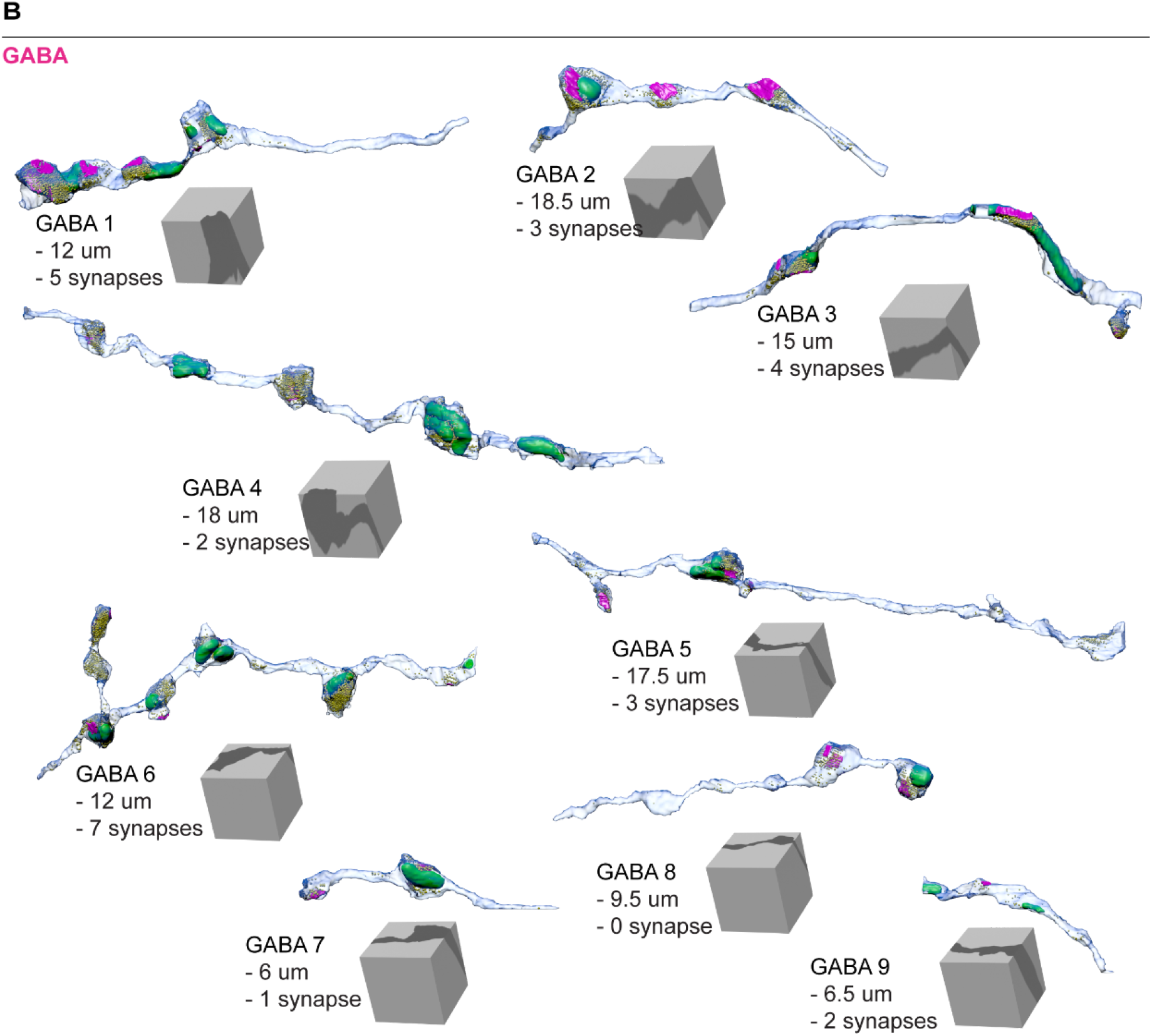
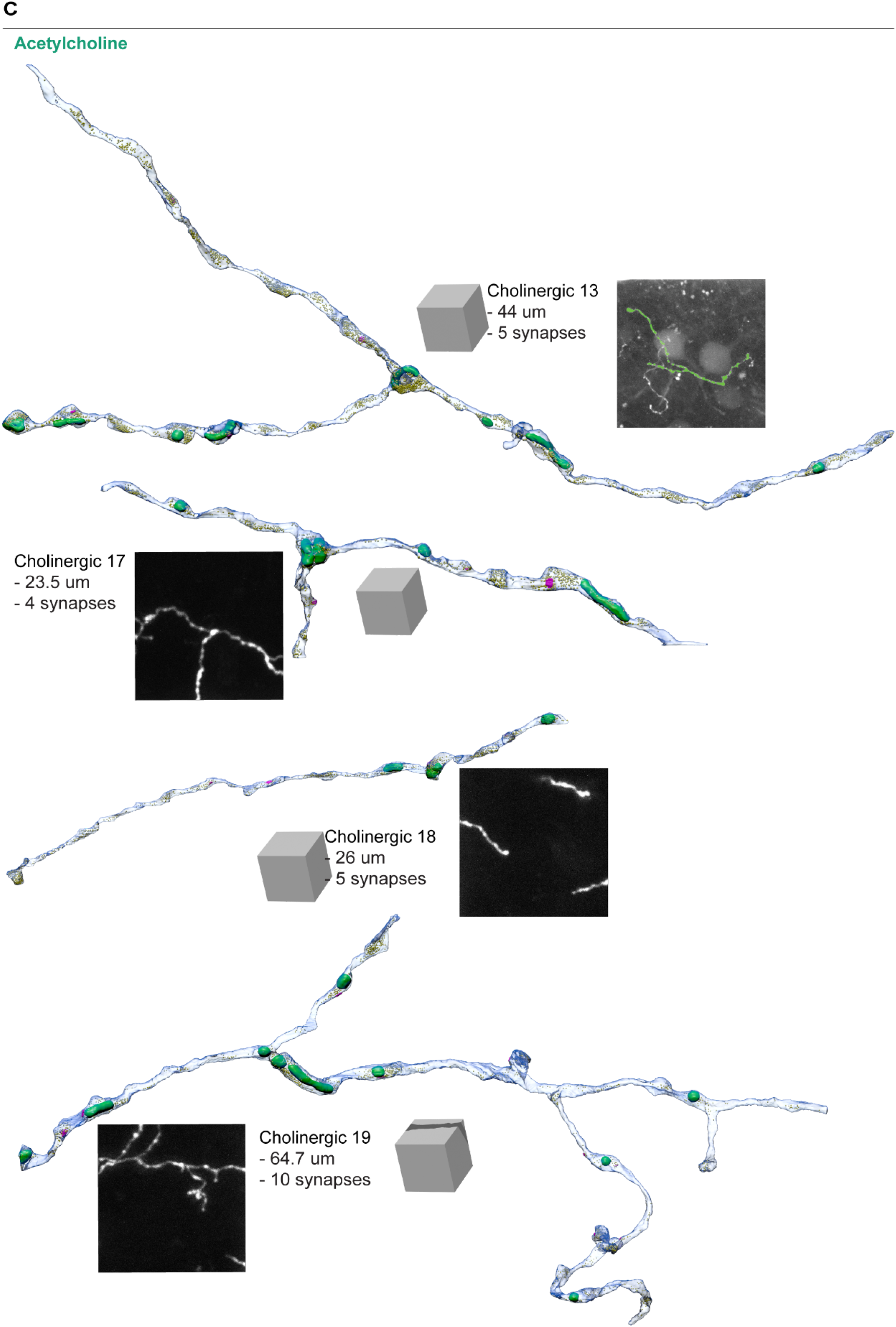
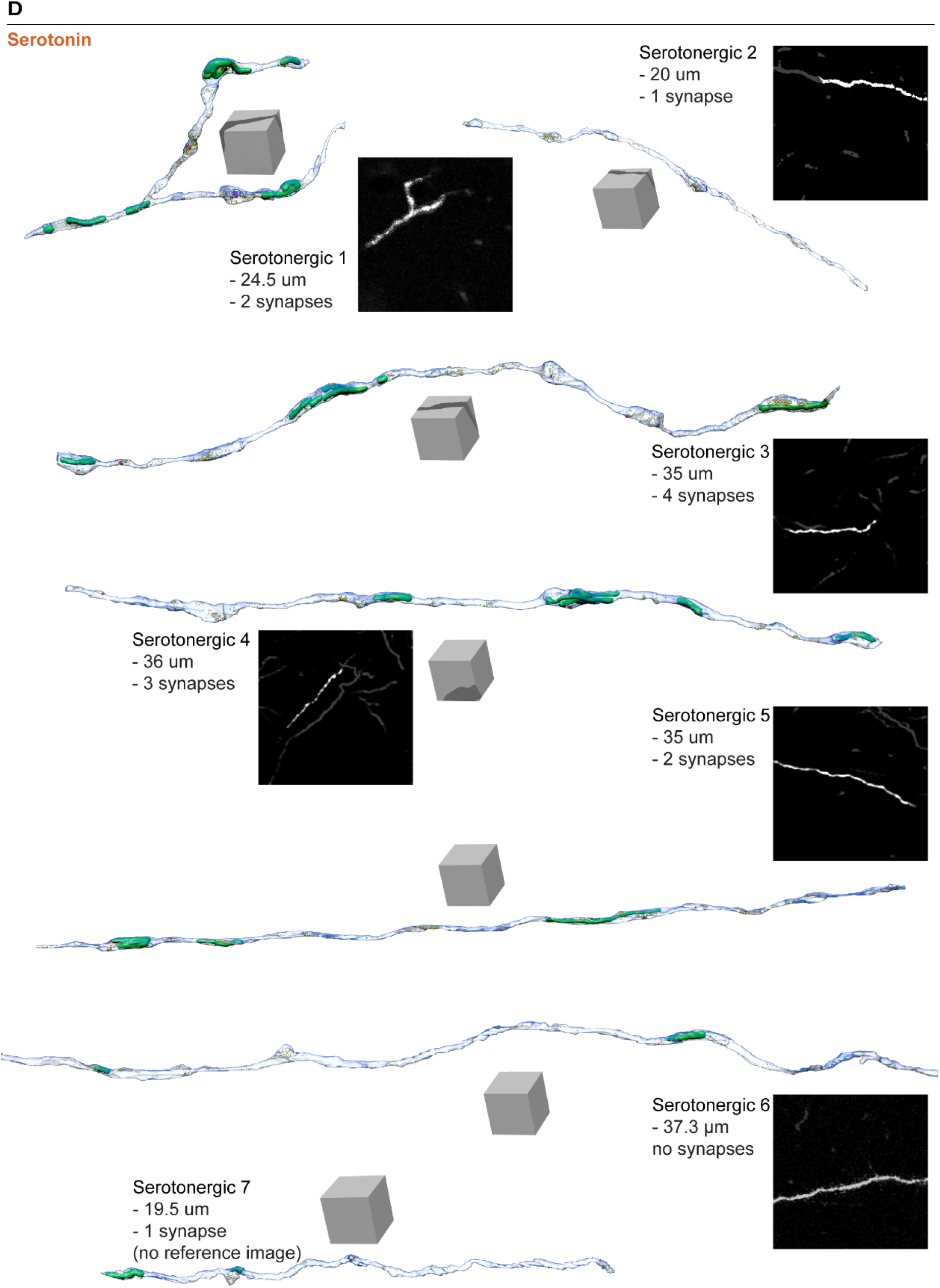
Best projections of axons from LM and Axon 3D reconstructions from FIBSEM. 3D model of reconstructed axonal segments for **(A)** Glutamate, **(B)** GABA, **(C)** Acetylcholine and **(D)** Serotonin respectively. For axons belonging to acetylcholine and serotonin, axons of interest were viewed with fluorescent microscopy (where present) and selected for FIBSEM imaging and reconstructed in 3D. Their total length, as well as the total number of synapses identified are indicated.

**Figure 2—figure supplement 1.**
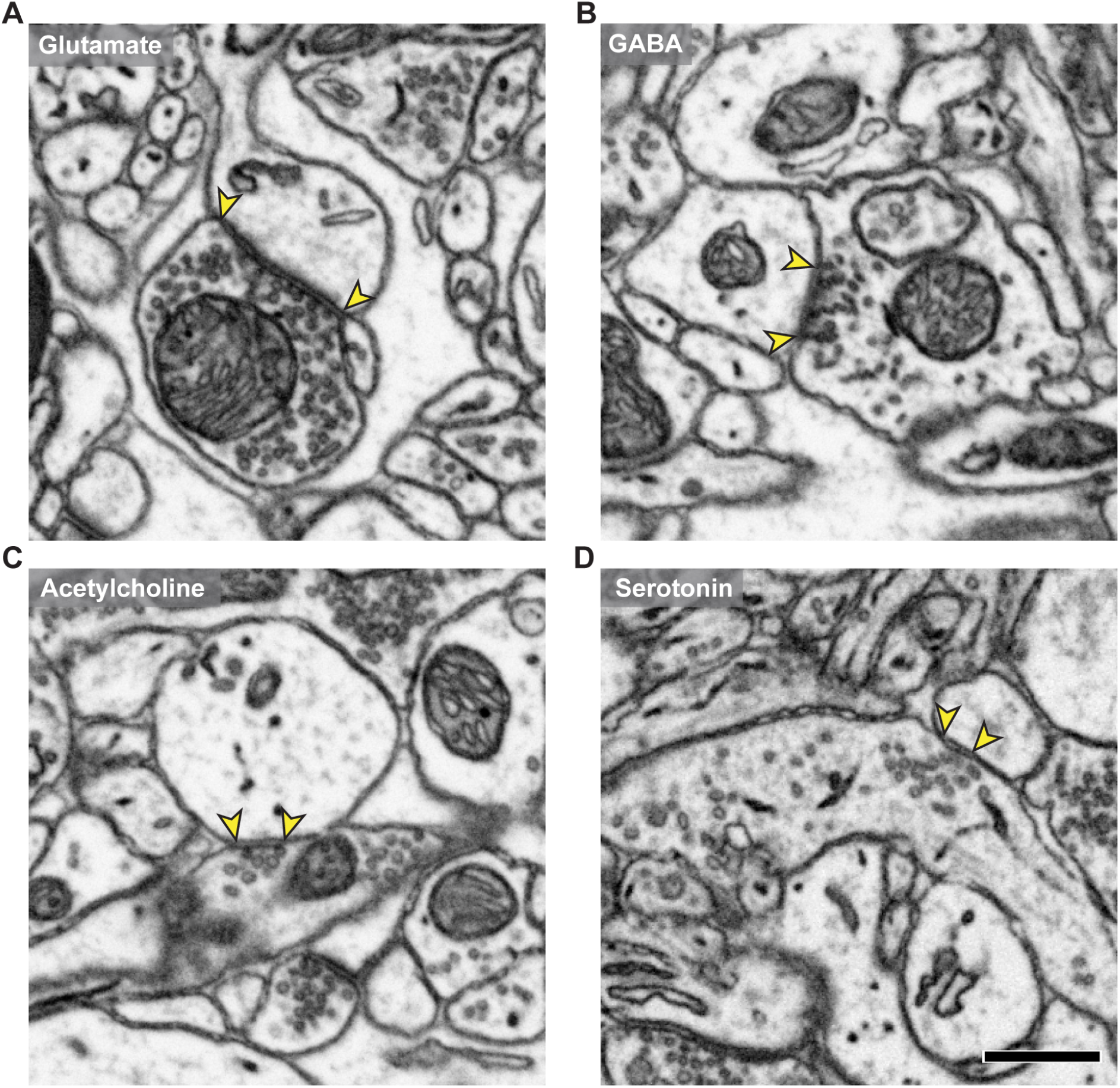
Electron micrograph of pre-postysnaptic membrane delimited and marked as synapses for each of the neurotransmitter subtypes. Representative electron micrographs of axonal varicosities of **(A)** glutamate, **(B)** GABA, **(C)** acetylcholine and **(D)** serotonin axons, containing vesicles within the varicosities, with arrows (yellow) indicating the synapses. Scale bar bottom right = 500 nm.

**Figure 3—figure supplement 1.**
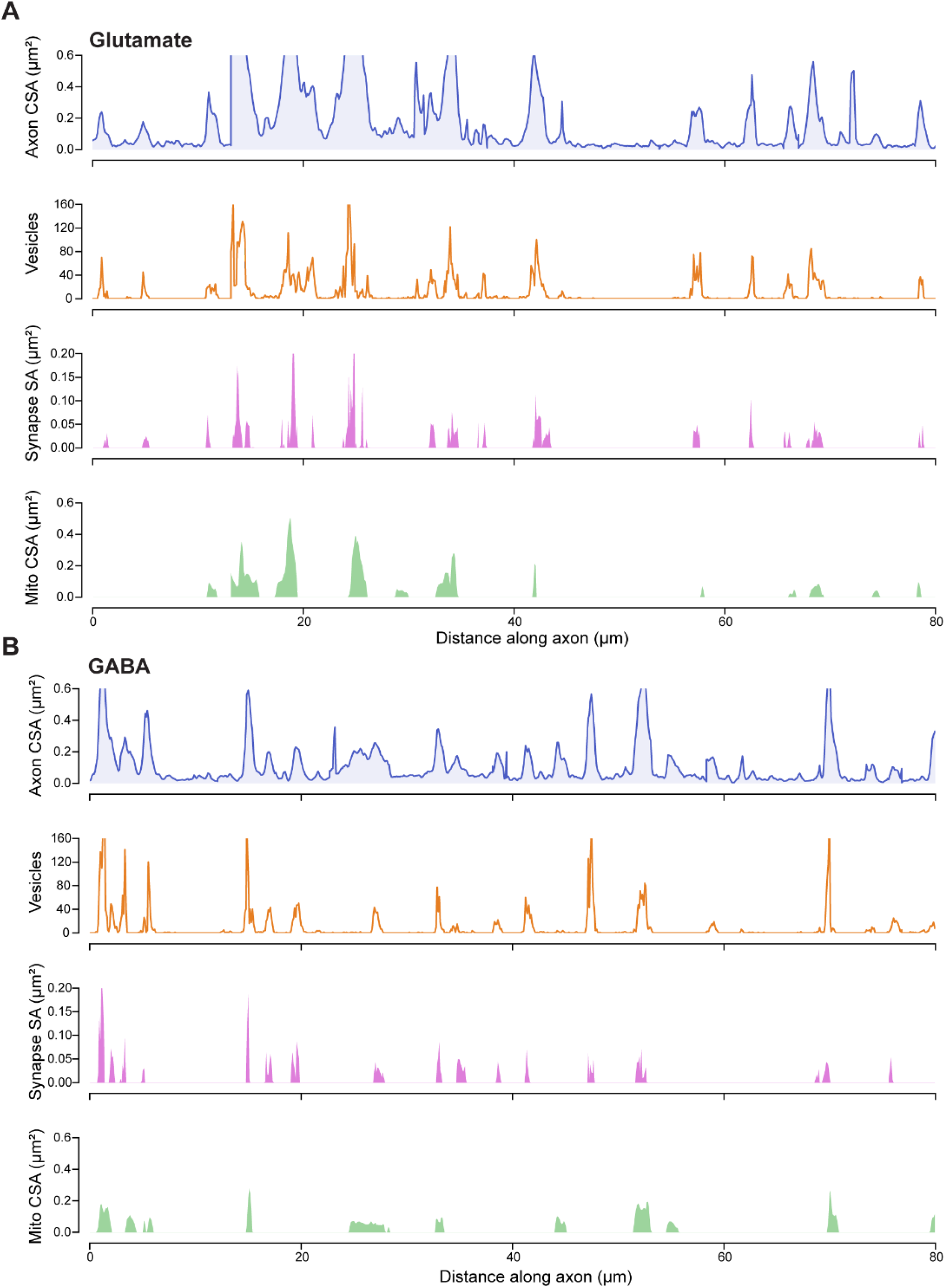
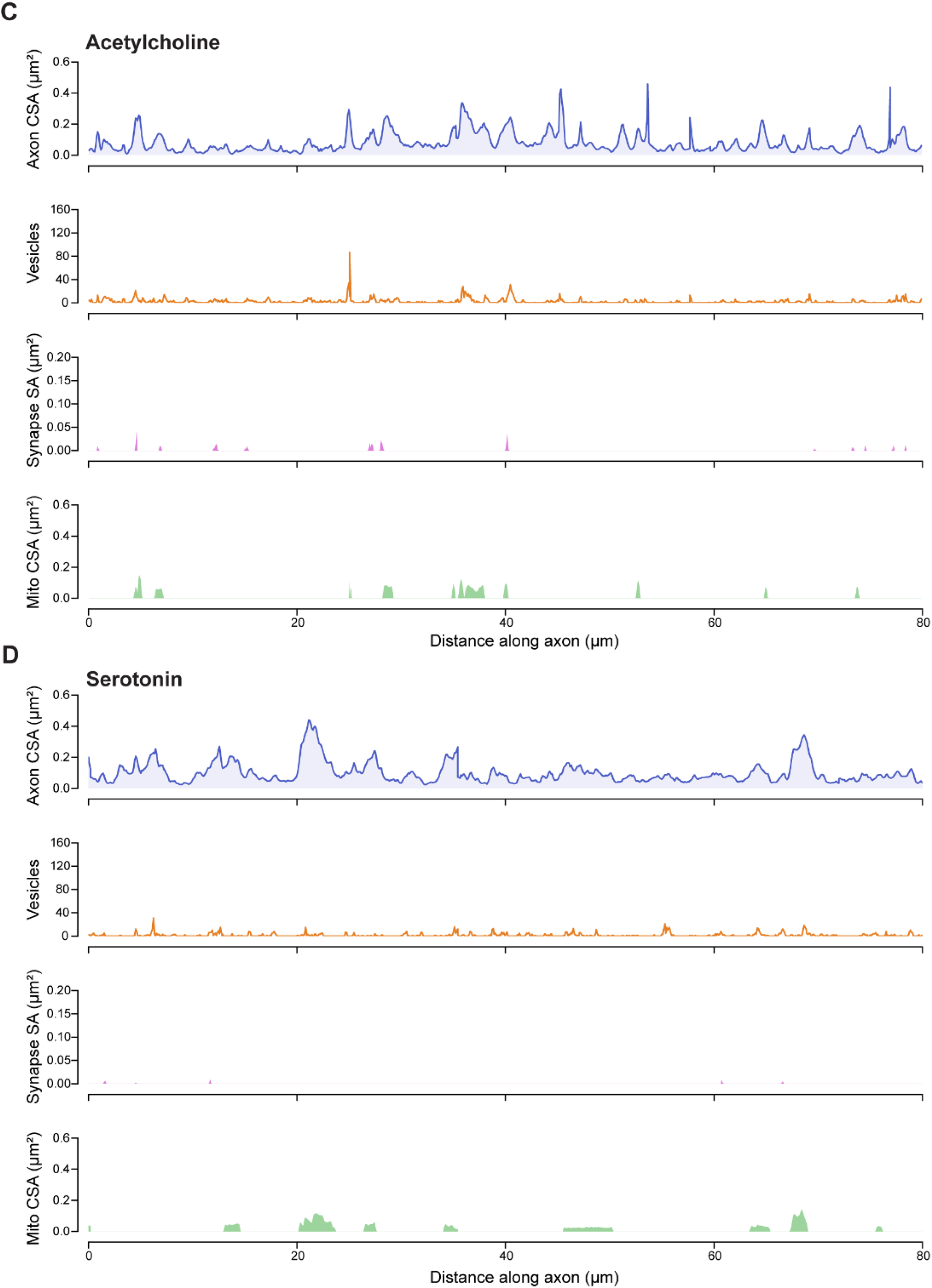
Expanded axon profile plots for each of the neurotransmitter subtypes shown in Figure 3. **(A)** Glutamate, **(B)** GABA, **(C)** Acetylcholine and **(D)** Serotonin respectively. From top to bottom: cross-sectional area (CSA) of the axon (blue), number of vesicles (orange), surface area (SA) of the synaptic contacts (pink) and cross-sectional area of mitochondria (green) along a length of axon (represented as distance along axon over 80 µm). All cross-sectional areas and surface areas are in µm^2^.

**Figure 4—figure supplement 1.**
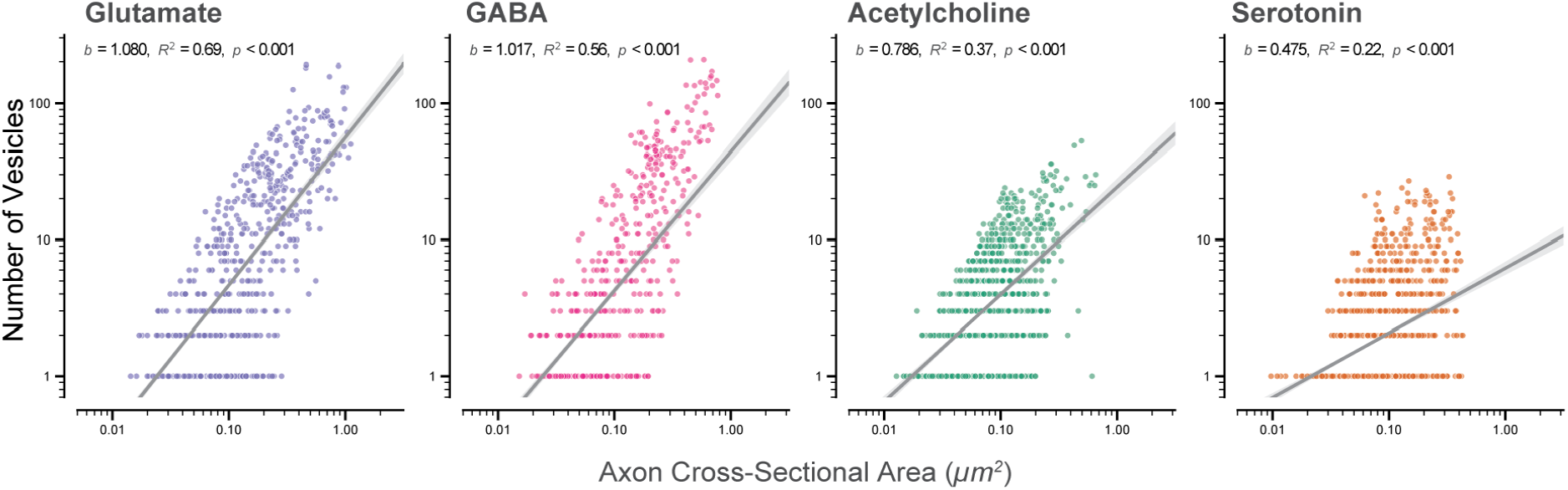
Allometric scaling relationship between the number of vesicles and cross-sectional area across neurotransmitter subtypes. Log-log regression plots (from left to right) showing the power-law relationship between the Number of Vesicles (counts, *y-axis*) and their corresponding axon cross-sectional area (*µm^2^*, *x-axis*) for glutamate, GABA, acetylcholine, and serotonin axon segments . Each data point represents an individual 100 nm axon segment. Solid grey lines indicate the best-fit linear regression on the log_10_-transformed data, and shaded regions represent the 95% confidence intervals. The scaling exponent (*b*), coefficient of determination (*R^2^*), and *p*-values are inset for each panel. Glutamatergic (*b* = 1.080, *R^2^* = 0.69, *n* = 1187) and GABAergic (*b* = 1.017, *R^2^* = 0.56, *n* = 1058) axons exhibit near-isometric scaling (*b* ≈ 1.0). In contrast, modulatory cholinergic (*b* = 0.786, *R^2^*= 0.37, *n* = 1646) and serotonergic (*b* = 0.475, *R^2^* = 0.22, *n* = 2076) axons display negative allometric scaling (*b* < 0.8), showing a weaker relationship between of the number of vesicles and the size of the axon. All slopes are significantly different from zero (*p* < 0.001).

**Figure 7—figure supplement 1.**
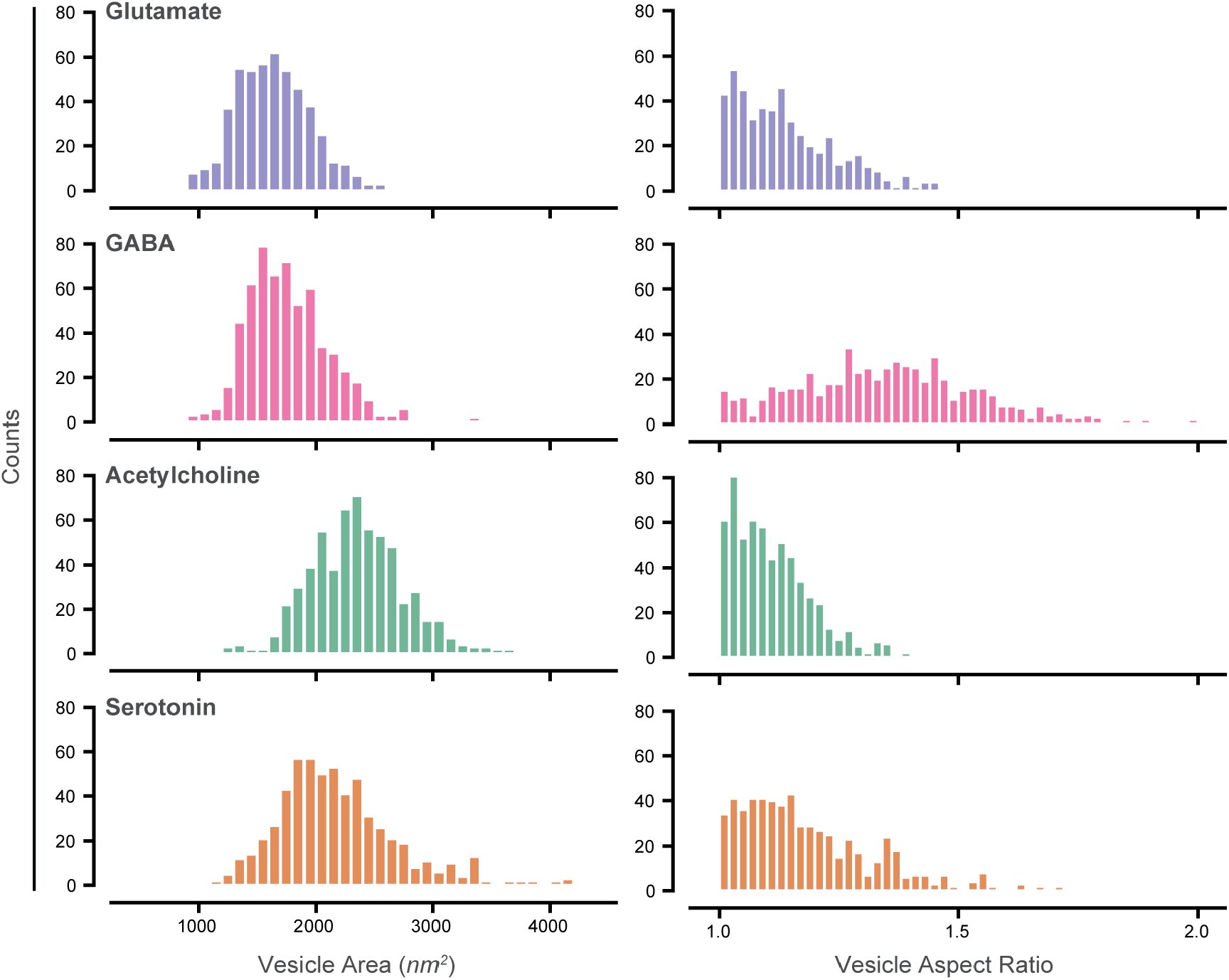
Distribution plot of vesicle area and aspect ratio. (left) Distributions of vesicle area across neurotransmitter subtypes. Histograms showing the frequency distribution of vesicle area measurements (nm^2^) in discrete bins of 100 nm^2^. From top to bottom, Glutamate, GABA, Acetylcholine and Serotonin. Vesicle area for glutamate and GABA appears to be tightly constrained while vesicles of neuromodulatory vesicles are larger and appear to have a wider spread. Note the right-skewed tail extending past 4000 nm^2^ within the distribution of Serotonin vesicles. (right) Distribution of vesicle aspect ratio across neurotransmitter subtypes. Histograms showing the frequency distribution of aspect ratio measurements in discrete bin width of 0.02. The distribution of Glutamate, Acetylcholine and Serotonin exhibits a sharp, highly concentrated peak near a baseline aspect ratio of 1.0, while the distribution for GABA is noticeably broader and flatter, extending more prominently toward higher aspect ratios (up to 2.0). The y-axes denote absolute count of vesicles within each bin (scaled from 0 to 80).

**Figure 8—figure supplement 1.**
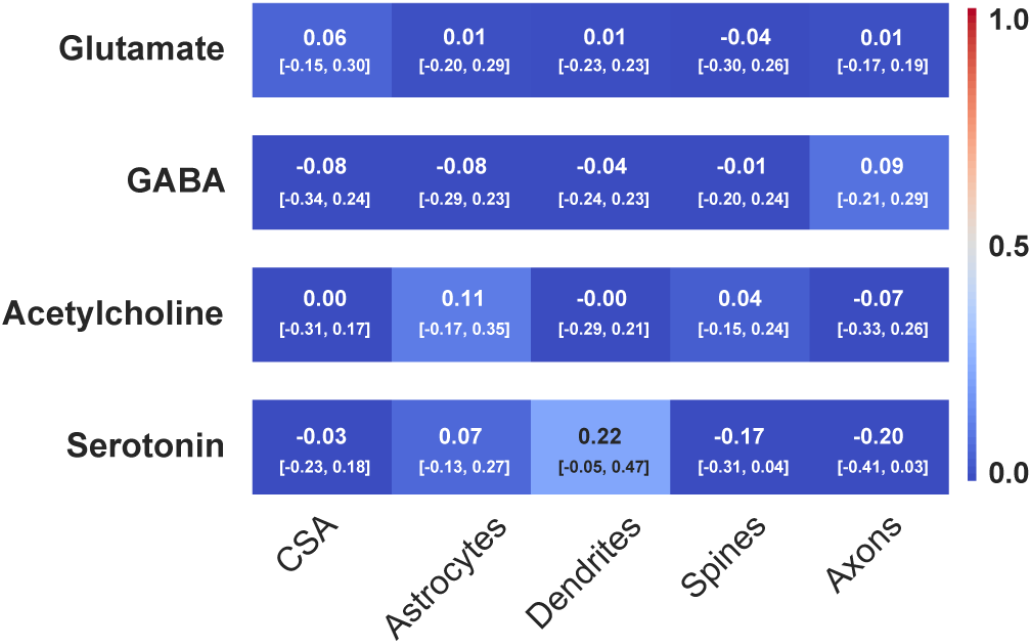
Null model for correlation specificity. Heat maps displaying correlation coefficients (*r*) for permuted vesicle counts against structure features of a simulated axon segment. In order to disrupt the biological pairings while preserving empirical frequency distributions, the vesicle numbers for the respective neurotransmitter subtypes: glutamate, GABA, acetylcholine and serotonin, were randomly shuffled across a simulated axon segment. Correlations were calculated against the axon cross-sectional area (CSA) and membrane contact areas: astrocytes, dendrites, spines and other axons. Colour mapping follows the scale bar (right), warmer colours indicate stronger positive correlations, whereas cooler colours indicate weaker correlations, as shown by the scale bar. Values within each cell indicate the observed *r*-value above its corresponding 95% bootstrap confidence interval [brackets]. Statistical significance was determined via label-permutation test. Metrics for both bootstrap and permutation were resampled over 10,000 iterations. All confidence intervals overlap zero, demonstrating that the pipeline is free of artificial correlation artifacts.

## Notes

### Competing Interest Statement

The authors have declared no competing interest.

